# Mucilage Produced by Sorghum (*Sorghum bicolor*) Aerial Roots Hosts Diazotrophs that Provide Significant Amounts of Nitrogen to the Plant

**DOI:** 10.1101/2023.08.05.552127

**Authors:** Rafael E. Venado, Jennifer Wilker, Vania Pankievicz, Valentina Infante, April MacIntyre, Emily S. A. Wolf, Saddie Vela, Fletcher Robbins, Paulo Ivan Fernandes-Júnior, Wilfred Vermerris, Jean-Michel Ané

## Abstract

Sorghum (*Sorghum bicolor*) is a significant crop globally, serving as an important source of food, feed, and fodder, and is increasingly recognized as an energy crop due to its high potential for biomass production. Certain sorghum accessions exhibit prolific aerial root development and produce abundant carbohydrate-rich mucilage after precipitation. This aerial root mucilage bears resemblance to that found in landraces of maize (*Zea mays*) from southern Mexico, which have previously been found to harbor diazotrophs. In this study, we examined the aerial root development of specific sorghum accessions and investigated the influence of humidity on this trait. Our microbiome analysis of the aerial root mucilage of maize and sorghum revealed the presence of numerous diazotrophs in sorghum mucilage, with *Pseudomonadota*, *Bacillota*, and *Bacteriodota* being the predominant phyla observed. However, the community composition varied significantly depending on the host plant and location. Through acetylene reduction, ^15^N_2_ gas feeding, and ^15^N isotope dilution assays, we determined that these sorghum accessions can acquire approximately 40% of their nitrogen from the atmosphere through these symbiotic associations on aerial roots. The nitrogen fixation occurring in sorghum aerial root mucilage presents a promising opportunity to reduce reliance on synthetic fertilizers and advance sustainable agricultural practices for food, feed, fodder, and bioenergy production.

## Introduction

Nitrogen (N) is an essential nutrient for plant growth and development. However, despite its abundance in the atmosphere as dinitrogen (N_2_), plants cannot access it directly [1]. Biological nitrogen fixation is the conversion of dinitrogen into ammonium performed by bacteria and archaea known as diazotrophs that express the nitrogenase enzyme [2,3]. This process is essential in agriculture, enabling certain plants, such as legumes, that associate with diazotrophs called rhizobia, to obtain large amounts of nitrogen directly from the atmosphere. Legumes host rhizobia in specialized organs called root nodules, inside which rhizobia fix nitrogen and transfer it to the plant in exchange for carbon from photosynthesis [4]. This symbiosis is critical for the productivity and sustainability of agricultural systems worldwide by reducing the need for synthetic fertilizers and enhancing soil fertility [5–7]. In contrast, cereal crops like maize (*Zea mays* L.) and sorghum (*Sorghum bicolor* (L.) Moench) generally require substantial nitrogen inputs. While synthetic nitrogen fertilizers, derived from natural gas using the Haber-Bosch process, can address this limitation, their extensive use has led to significant economic and environmental concerns, including nitrogen leaching and the generation of greenhouse gases. For decades, approaches have been proposed to enhance biological nitrogen fixation in cereals, including engineering root nodules in cereals or the expression of the nitrogenase directly in plant cells [8,9]. Another strategy is to manipulate the diazotrophs naturally associated with cereals. Such associative diazotrophs have been documented, for instance, in the rhizosphere, the xylem, or the root mucilage of cereals [10,11]. A synthetic community comprising *Klebsiella variicola* MNAZ1050 and *Citrobacter sp* MNAZ1397 was reported recently to contribute 11% of nitrogen maize through biological nitrogen fixation [10]. Some approaches also rely on altering the genetic circuits controlling nitrogenase expression in associative diazotrophs to induce the release of ammonium on the root surface [12]. While exciting, all these engineering approaches require long development times and are often based on complex genetic manipulations that limit their development and adoption. To enhance the benefits of biological nitrogen fixation for cereal crops in the shorter term, we explored plant natural diversity to identify genotypes of cereals that could be better hosts for diazotrophs than currently available cultivars.

In 2018, we and colleagues published the characterization of maize landraces cultivated and selected by Indigenous communities in the Sierra Mixe, Oaxaca, Mexico, that secrete viscous mucilage from their aerial roots. Unlike brace roots, these aerial roots are adventitious nodal roots that do not reach the ground. We demonstrated that this mucilage supports communities of diazotrophs that can provide 29-82% of the plant’s nitrogen needs [13–16]. These landraces could serve as the foundation for nitrogen fixation in maize through conventional breeding with limited impact on yield [17]. However, the production of aerial roots is not unique to maize. Such roots have been reported throughout the Andropogoneae and Paniceae tribes that include maize, sorghum, sugarcane (*Saccharum* spp.), foxtail millet (*Setaria italica* L.), and pearl millet (*Pennisetum glaucum* (L) R. Br.). These roots serve anchorage functions, especially under flooded conditions, and are important for water and nutrient acquisition [18,19]. Sorghum is a versatile and resilient crop used for food, feed, and fodder production. It has recently gained significant attention as a promising bioenergy crop due to the high biomass productivity of some genotypes [20,21]. Its ability to thrive in diverse climatic conditions, including drought-prone regions, makes it an attractive option for sustainable food and bioenergy production worldwide [22,23].

Despite the similarity between sorghum aerial roots and those in maize known to host diazotrophs, evidence of biological nitrogen fixation in sorghum has yet to be well characterized. Images of aerial roots of sorghum covered with mucilage, as well as a micrograph depicting the bacterium *Azospirillum brasilense,* were featured in the abstract book of a conference on nitrogen fixation in cereals hosted by the International Crops Research Institute for the Semi-Arid Tropics, India (ICRISAT) in 1984. Despite this decades-old report, no data have been published on biological nitrogen fixation in sorghum aerial roots [24]. The similarity between these sorghum aerial roots and those from the Sierra Mixe maize prompted us to explore nitrogen fixation in sorghum. We examined the diversity of aerial roots in sorghum accessions, mucilage production, and their microbiome. We demonstrated that, like the maize landraces from the Sierra Mixe, some sorghum accessions can acquire significant amounts of nitrogen from their aerial roots via the diazotrophs established in their mucilage.

## Results

### Specific sorghum accessions developed aerial roots

Sorghum accessions selected from the sorghum minicore [25] and augmented with accessions provided by ICRISAT exhibited extensive development of thick aerial roots, similar to those in Sierra Mixe maize landraces [13,25]. Fourteen accessions were grown in a greenhouse at the University of Wisconsin-Madison and assessed for the number of nodes with aerial roots to characterize this trait better (**Supplementary Table 1**). The number of nodes with aerial roots varied from 0 to 8 between accessions. Accessions IS17348 and IS24453 exhibited minimal and no development of aerial roots, respectively. (**Figures 1A**). Upon exposure to water, a number of accessions demonstrated substantial mucilage production from their aerial roots. Notably, accessions IS10757, IS11026, IS15170, IS15466, IS23992, IS25089, and IS28025 consistently produced an average of 200 µl of mucilage per aerial root with comparable lengths. However, accessions IS17348, IS24453, IS28023, IS28033, IS28262, and IS29078 exhibited lower mucilage production, averaging less than 100 µl per root with similar lengths. (**Figure 1B and C**). Sorghum aerial root diameters were measured to determine if the sorghum accessions exhibited thick aerial roots, as shown for the Sierra Mixe maize landraces. The diameter of aerial roots ranged from 0 (when absent, only the accession IS24453) to 8 mm (**Figures 1D and 1E**). Through this screening, we identified IS10757, IS11026, IS15170, and IS25089 as the most promising accessions, distinguished by both thick aerial roots and high mucilage production. These results indicate significant variation in the number and diameter of aerial roots and mucilage production among sorghum accessions. A positive correlation was observed between the number of nodes forming aerial roots, root diameter, and mucilage production volume (**Supplementary Figure 1A**). The correlation between the aerial root diameter and mucilage volume was modest, r^2^ = 0.4 (p-value = 0.0024), but low for the number of nodes producing aerial roots and volume, r^2^ = 0.2 (p-value = 0.1517, non-significant) (**Supplementary Figure 1B**). This result indicates that, as we reported for maize, thicker roots are associated with higher mucilage production in these sorghum accessions, too [15].

**Fig 1.**
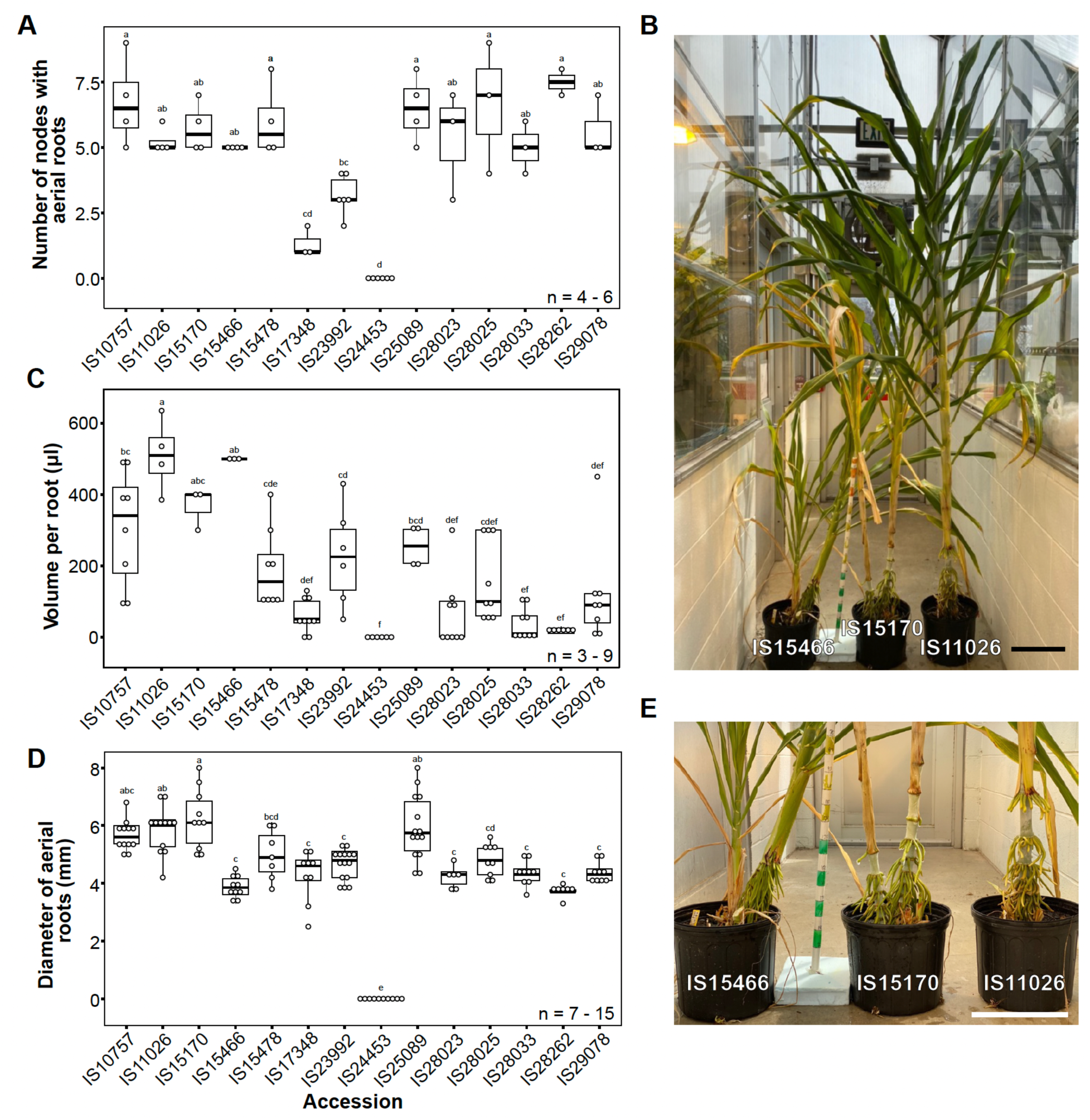
Sorghum screening of selected accessions under greenhouse conditions. Different sorghum accessions display diverse traits associated with aerial roots and mucilage production. A) Number of nodes in different sorghum accessions (n = 3-6). Most of the accessions evaluated showed more than 5 nodes with aerial roots. B) Representative image of selected sorghum accessions. Scale bar = 28 cm. C) Volume of mucilage per root (μl) produced by different sorghum accessions (n = 4-9). The production of mucilage varied among all the accessions. D) Diameter (mm) of aerial roots in different sorghum accessions (n = 6-15). Thick roots with diameters bigger than 4 mm were noted in nearly all the accessions. E) Aerial roots of the sorghum accessions in D. Scale bar = 28 cm. An ANOVA (F-test) to calculate significant differences between means was conducted with the package multcompView (0.1-10).

To test these findings under field conditions, the same set of 14 sorghum accessions were grown at the University of Florida North Florida Research and Education Center-Suwannee Valley in 2021 (Live Oak, Florida, USA) and the University of Wisconsin-Madison West Madison Agricultural Research Station in 2022 (Madison, Wisconsin, USA). Florida has a humid subtropical climate, whereas Wisconsin has a continental climate that is affected by the humidity from Lake Superior and Lake Michigan. In these field experiments, the number of nodes with aerial roots, the root diameter, and the number of roots at the top node were recorded (**Supplementary Figure 2**). Environmental conditions, likely a combination of soil properties and humidity, played a significant role, as not all accessions developed aerial roots in both locations. The sorghum accessions generally produced more aerial roots in Florida than in Wisconsin. In Florida, approximately half of the 14 accessions produced nodes with aerial roots; in Wisconsin, fewer than a quarter of the accessions did (**Supplementary Figures 2A and B**). Furthermore, larger aerial root diameters were recorded in Florida compared to Wisconsin (**Supplementary Figures 2C and D**). In Florida, the count of aerial roots at the top node exceeded that in Wisconsin, averaging six roots compared to three roots at the top node, respectively (**Supplementary Figures 2E and F**). Overall, accessions IS11026, IS15170, and IS23992 demonstrated the potential to develop traits related to aerial root formation and mucilage production, highlighting the significant influence of environmental factors on these characteristics.

### Humidity influences the number of nodes in sorghum

Environmental factors can affect plant growth and development, including moisture, light intensity, and temperature [26]. We investigated the impact of humidity on our previous findings under greenhouse and field conditions,1 we evaluated two sorghum accessions: IS23992, which produces many aerial roots, and IS24453, which lacks aerial roots under field conditions (**Figure 1 and Supplementary Figure 2**). We grew these accessions in a greenhouse under either high humidity (75%) or low humidity (30%) conditions. For both sorghum accessions, we observed a significant increase in the number of nodes with aerial roots when grown under high humidity, with a more pronounced effect observed in IS23992 (**Figure 2A**). Interestingly, IS24453 exhibited aerial root development in this experiment, contrasting with the results observed in the field experiments (**Supplementary Figure 2**). Moreover, the number of aerial roots at the top node was only significantly affected by humidity in IS24453 (**Figure 2B**). To determine whether the increase in the number of nodes with aerial roots in humid conditions was due to differences in root diameter, we measured the diameter of aerial roots at the top node under both humidity conditions. The aerial root diameter did not significantly differ between these accessions (**Figure 2C**). These findings highlight the critical role of humidity as a key environmental factor in aerial root development in sorghum.

**Fig 2.**
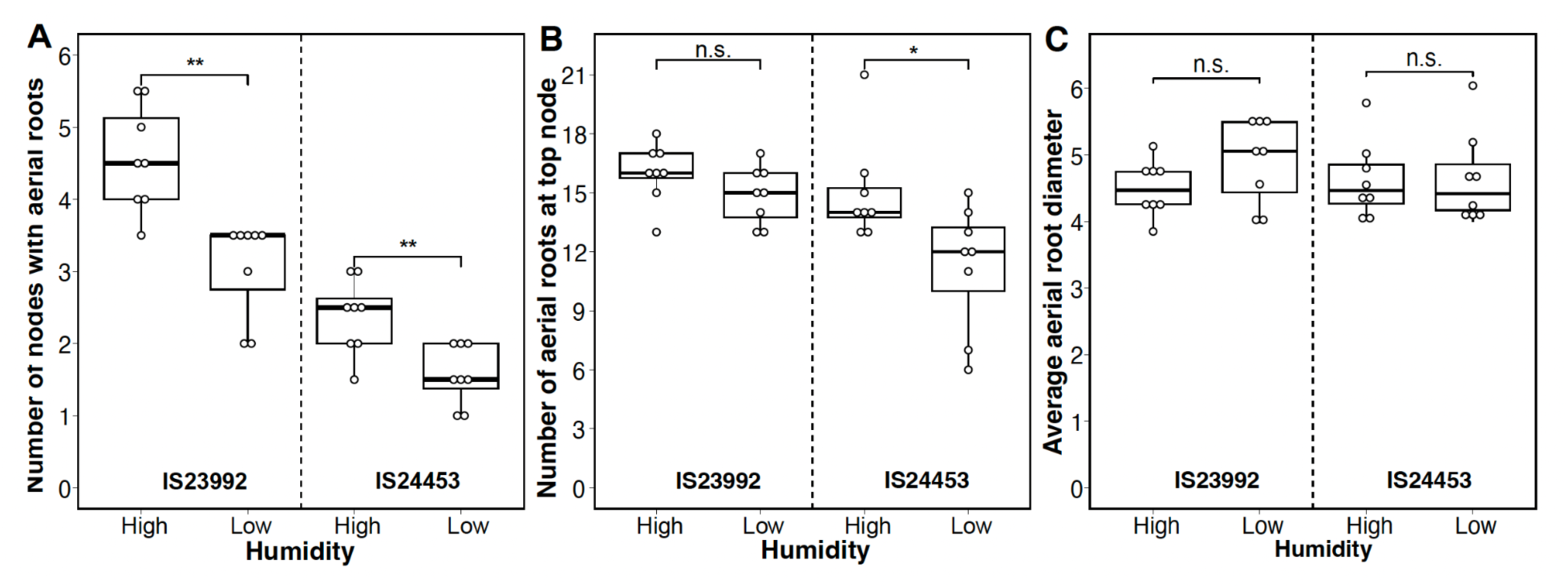
Effect of humidity in sorghum accession IS23992 and IS24453. Boxplots of different phenotypes measured in the sorghum accessions under high (75%) and low (30%) relative humidity. A) Number of nodes with aerial roots (n = 8). The number of nodes with aerial roots significantly increased in both accessions. B) Number of roots on the top node (n = 8). Only IS24453 increased the number of roots at the top node. C) Average root diameter on the top node (n = 8). The root diameter was not affected by the humidity in both accessions. A Wilcoxon test was performed in R (Ver 4.2.1) with the package ggpubr (Ver 0.6.0). Significance levels * p-value ≤ 0.05, ** p-value ≤ 0.01, and n.s. not significant.

### Sorghum mucilage is composed of carbohydrates

The viscous mucilage produced by the Sierra Mixe maize landraces consists of a complex polysaccharide with a galactose backbone and side chains of fucose, arabinose, xylose, mannose, and glucuronic acid [13,14]. Visually the mucilage from different sorghum accessions, such as IS2392 (**Figure 3A**), looked similar. To identify any chemical differences between maize and sorghum mucilage, we conducted a total carbohydrate composition analysis using gas chromatography/mass spectrometry (GC/MS). We collected mucilage from accession IS11026 in the greenhouse, as this accession produced a substantial amount of mucilage under greenhouse conditions (**Figure 1C**). For a clear distinction between sorghum and maize mucilage comparison, we presented the proportions (mole %) of the quantified monosaccharides. (**Figure 3B and Supplementary Table 2**). Arabinose, galactose, glucuronic acid, and mannose had higher ratios in sorghum than in maize mucilage. In particular, galactose accounts for nearly 50% of the total sugars in sorghum mucilage, and the proportion of fucose in sorghum mucilage was approximately half of that in maize mucilage. Using this approach, we did not detect the presence of other carbohydrates, such as glucose, rhamnose, ribose, and galacturonic acid. These results indicate that the aerial root mucilage of the two species consists of the same monosaccharides, but that their proportions differ.

**Fig 3.**
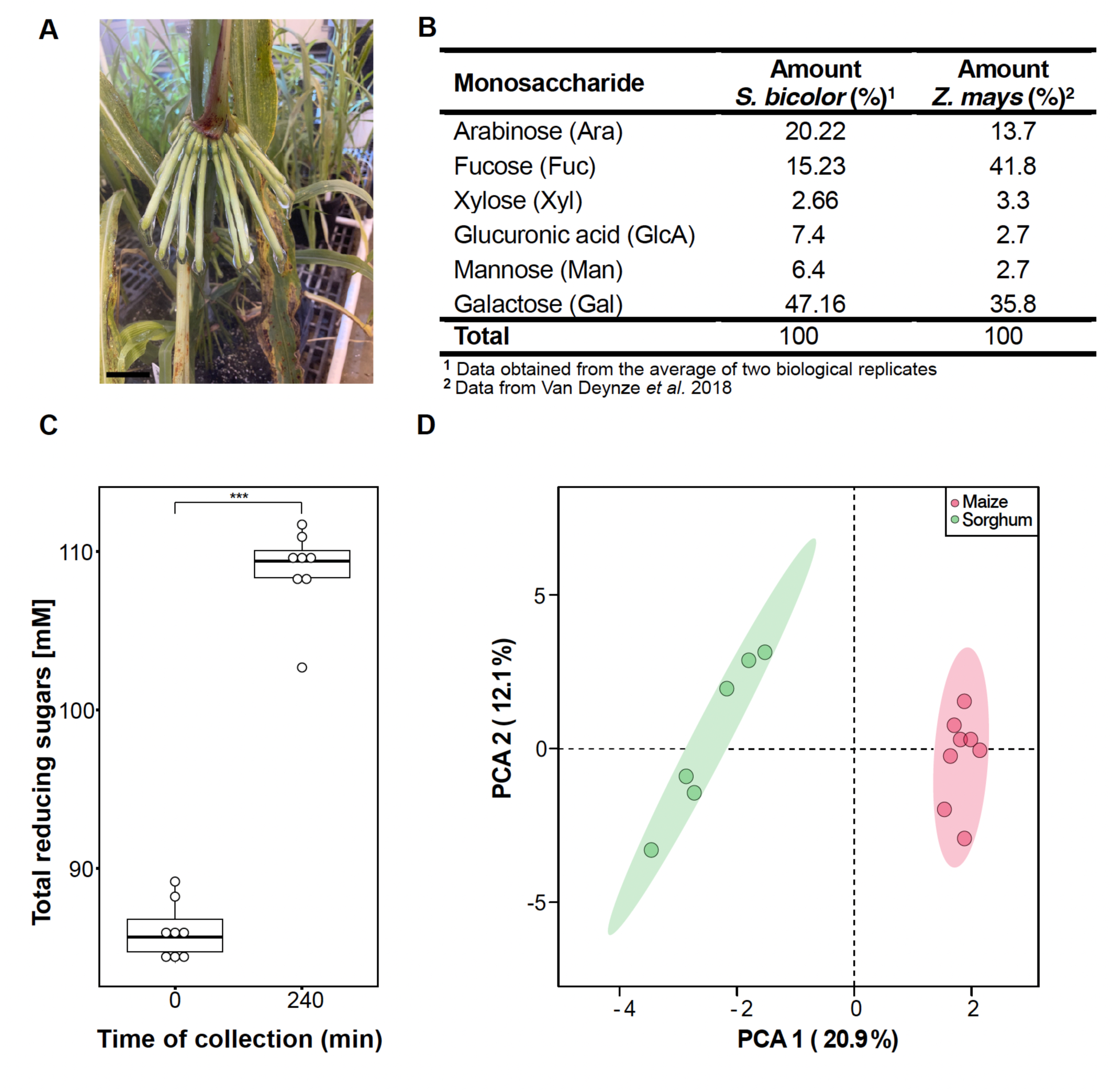
Sorghum aerial root mucilage composition. Sorghum aerial roots release mucilage. A) Mucilage production is triggered by water and under high-humidity conditions. The mucilage from different accessions appears similar. Accession IS2392 serves as a representative example of sorghum mucilage. Scale bar (5 cm) B) Monosaccharide composition of sorghum accession IS11026 and Sierra Mixe maize mucilage. Both mucilages consist of the same monosaccharides but in different proportions (mole %). C) Concentration of reducing sugars in sorghum mucilage from the accession IS23992 determined with benedict’s reaction. A Wilcoxon test was performed to compare means *** p-value ≤ 0.001 (n = 8). D) Principal component analysis using Sparse Partial Least Squares. Two clusters are depicted based on the species studied: sorghum accessions IS29091 and IS2245 and maize landraces CB017456 and GRIN 19897.

In maize landraces from the Sierra Mixe, water triggered the expression of plant genes associated with carbohydrate synthesis and degradation [15]. To investigate this degradation process further, we assessed the levels of free sugars in sorghum mucilage at 1 and 5 hours after adding water to the aerial roots of the accession IS23992. Using Benedict’s reaction to quantify free sugars, we observed that the free sugar concentration increased over time, rising from an average of 86 mM at 1 hour to 109 mM after 5 hours (**Figure 3C**)[27]. This finding suggests that sorghum aerial roots produce enzymes that break down the mucilage polysaccharide into simple sugars. We cannot exclude the contribution of microbes in this degradation process as the aerial roots from plants grown in the greenhouse were not germ-free. However, compared to field conditions, very few microbes could be isolated from sorghum mucilage when grown in this greenhouse.

To explore the differences in the metabolites between maize and sorghum aerial root mucilage, we performed a metabolomics analysis using gas chromatography coupled with time-of-flight mass spectrometry (GC/TOF-MS) (**Supplementary Table 3**). We utilized the sorghum accessions IS29091 and IS2245 from the Wolf *et al.* (2023) [28] study, which investigated the diversity of aerial roots in sorghum, along with maize landraces CB017456 and GRIN19897, which display aerial root. In total, 252 metabolites were detected in maize and sorghum mucilage (**Dataset S1)**. A principal component analysis (PCA) revealed two clusters where samples were grouped based on plant species (**Figure 3D**). We used a Significance Analysis of Microarray (SAM) to identify metabolites that differed among the mucilage samples of the two species. The top 10 most differentially abundant metabolites between sorghum and maize included 6 unknown metabolites but also octadecanoic acid, montanic acid (octacosanoic acid), pinitol (1D-*chiro*-inositol), and *N*-acetyl aspartate diethyl ester (**Supplementary Table 4**). These metabolomic differences may contribute to functional differences between sorghum and maize mucilage.

### Aerial root mucilage from sorghum contains abundant and large border cells

The root cap of underground roots produces small quantities of mucilage [29–31]. In particular, border cells (BCs) that detach from the root cap are responsible for mucilage production [15,32]. To investigate sorghum mucilage production, we examined mucilage derived from both underground and aerial roots in sorghum accession IS23992 (**Supplementary Figures 3A and C**). BCs derived from underground roots exhibited a distinct and elongated morphology similar to those observed in BCs from pea (*Pisum sativum* L.), maize, and alfalfa (*Medicago sativa* L.)[33–35]. In contrast, BCs from sorghum aerial roots appeared much larger and elongated, with irregular shapes and sizes (**Supplementary Figure 3C**). Interestingly, these BCs from sorghum aerial roots resembled those recently reported in mucilage from maize aerial roots [15]. These larger BCs may contribute to the abundant mucilage production by maize and sorghum aerial roots.

Border cells detached from the underground root cap can remain active for several weeks [32,36]. To determine the viability of sorghum border cells, we used a live-dead staining protocol on both types of sorghum mucilage collected after 72 hours. We used a combination of SYTO^TM^ 13 and propidium iodine, which are permeating and non-permeating cell dyes, respectively [37,38]. Most cells were viable, as indicated by only SYTO^TM^ 13 in the nuclei. Additionally, BCs from underground roots displayed regular and well-defined nuclei, while BCs from aerial roots exhibited irregular nuclei (**Supplementary Figures 3B and D**). To quantify the morphological differences between the two types of BCs, we measured their length between the two cellular poles and their cell area. Significant differences were observed in both measurements. BCs from aerial roots are generally twice the size of BCs from underground roots. Although both types of BCs maintain the function of mucilage production, the observed morphological differences may account for the variation in the amount of mucilage secreted by these cells.

### Sorghum mucilage hosts a wide range of diazotrophs

To explore the microbial diversity within sorghum mucilage, we isolated and identified bacteria using 16S amplicon sequencing. The sorghum association panel [39] and sorghum minicore [25] were used to explore the genetic control of aerial root development in sorghum [28], obtain large amounts of mucilage, and characterize aerial root microbiomes [28,39]. We used mucilage samples from accessions IS2245, IS23992, IS29092 and IS2902 from the fields at the University of Wisconsin West Madison Agricultural Research Station and the UF North Florida Research & Education Center - Suwannee Valley. We also included mucilage samples from Sierra Mixe landraces at a neighboring West Madison Agricultural Research Station field as reference (**Supplementary Table 5**). To analyze the bacterial composition, we amplified and sequenced the 16S V3-V4 region and grouped the reads into Operational Taxonomic Units (OTUs) to provide information on the relative abundance of different taxa [40]. We performed principal coordinate analysis to examine the similarity between samples. As expected, the plant origin of the mucilage was the main factor influencing beta diversity, although location also plays a role in this diversity (**Figure 4A**). However, Analysis of Molecular Variance (AMOVA) uncovered instances where no statistically significant differences were detected between the microbiomes of maize and sorghum (**Supplementary Table 6**). We also investigated the taxonomy of bacteria in sorghum and maize mucilage. The phylum Pseudomonadota (former Proteobacteria) was dominant in the mucilage of both maize and sorghum, followed by Bacillota and Bacteroidota (**Figure 4B**). The profiles of samples from Wisconsin were relatively similar regardless of the plant of origin, while samples from Florida exhibited enrichment in Cyanobacteria (**Figure 4B**). In both mucilage samples, 34 distinct bacterial genera were identified, showing, as expected, differences in their abundance across the samples (**Supplementary Figure 4)**.

**Fig 4.**
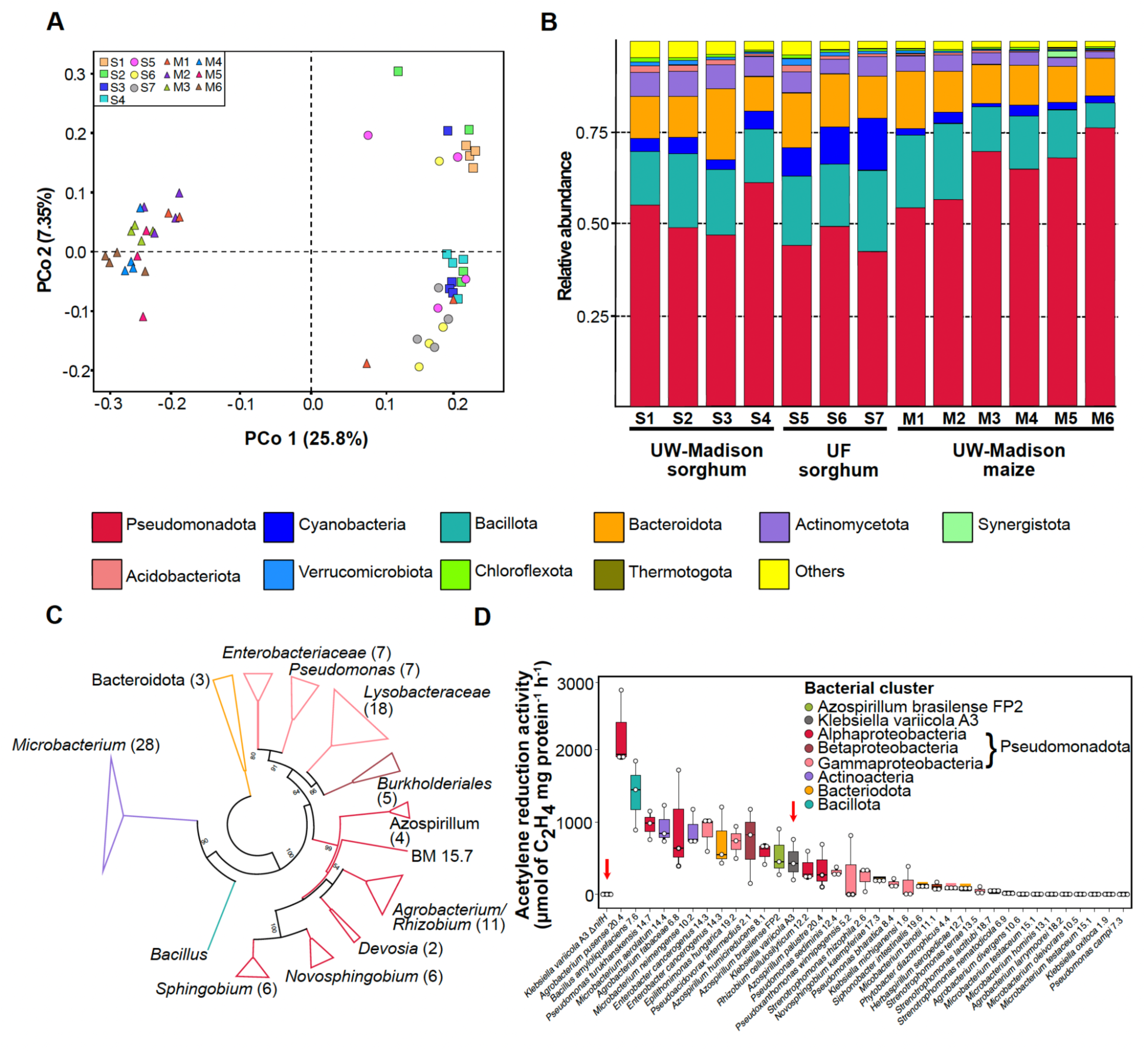
Microbiome profile of sorghum mucilage. The community profile shows the relative abundance of bacteria present in sorghum (S) and maize (M) mucilages. Samples S1 to S4 (squares) and M1 to M6 (triangles) were collected from the field at the University of Wisconsin-Madison (UW-Madison), and samples S5 to S7 (circles) were collected at the University of Florida (UF). A) Principal coordinates analysis plot of sorghum and maize samples based on 16S amplicon sequences. B) Relative abundance of sorghum and maize samples based on 16S amplicons and OTUs. C) Unrooted maximum-likelihood phylogenetic tree of 103 bacteria isolated from sorghum aerial roots mucilage in BMGM nitrogen-free semisolid medium. The evolutionary distances were computed using the Jukes-Cantor method and are in the units of the number of base substitutions per site. This analysis involved 103 nucleotide sequences. All positions containing gaps and missing data were eliminated (complete deletion). There were 908 positions in the final dataset. Numbers at nodes are bootstrap values (1000 replications, values < 50% are not shown). Colors: Red: Gammaproteobacteria, Purple: Betaproteobacteria, Blue: Alphaproteobacteria, Brown: Bacillota, Green: Actinomycetota, Yellow: Bacteroidota. D) Nitrogenase activity of 34 isolated from sorghum aerial roots mucilage in BMGM nitrogen-free semisolid medium (n=3). The red arrows show the reference strains for comparison. Bacteria: *Klebsiella variicola* A3 Δ*nifH*. Species codes (alphabetic order): *Agrobacterium divergens*, *Agrobacterium fabaceae*, *Agrobacterium larrymoorei*, *Agrobacterium pusense*, *Azospirillum brasilense*, *Azospirillum humicireducens*, *Azospirillum palustre*, *Bacillus amyloliquefaciens*, *Enterobacter cancerogenus*, *Epilithonimonas hungarica*, *Herbaspirillum seropedicae*, *Klebsiella michiganensis*, *Klebsiella oxytoca*, *Klebsiella variicola*, *Microbacterium aerolatum*, *Microbacterium binotii*, *Microbacterium hominis*, *Microbacterium neimengense*, *Microbacterium oleivorans*, *Microbacterium testaceum*, *Novosphingobium kaempferiae*, *Phytobacter diazotrophicus*, *Pseudoacidovorax intermedius*, *Pseudomonas bharatica*, *Pseudomonas campi*, *Pseudomonas lutea*, *Pseudomonas sediminis*, *Pseudomonas turukhanskensis*, *Pseudoxanthomonas winnipegensis*, *Rhizobium cellulosilyticum*, *Siphonobacter intestinalis*, *Stenotrophomonas lactitubi*, *Stenotrophomonas nematodicola*, *Stenotrophomonas rhizophila*, and *Stenotrophomonas terrae*.

In a second experiment, we isolated bacteria from the same sorghum association panel but expanded the number of accessions (**Supplementary Table 7**) [28,39]. The isolation of diazotrophs was performed using the nitrogen-free semi-solid BMGM medium, which revealed a high diversity of potential diazotrophs. We isolated 103 non-clonal strains from 34 different bacterial species with the prevalence of Pseudomonadota (Gammaproteobacteria - 31, Alphaproteobacteria - 33, Betaproteobacteria - 5), Actinomycetota (28), Bacteroidota (5), and Bacillota (1) (**Figure 4C and Supplementary Table 7**). Subsequently, we assessed the nitrogenase activity via the acetylene reduction assay (ARA) in one representative strain selected from our prior isolates. Additionally, we introduced the reference strains *Azospirillum brasilense* FP2[41], a bacterium known for promoting plant growth and commonly associated with crops, and *Klebsiella variicola* A3, isolated from sorghum mucilage and established as a standard for nitrogen fixation in our laboratory. As a negative control, we included the *Klebsiella variicola* A3 Δ*nifH* mutant, unable to fix nitrogen. The strains *Agrobacterium pusense* 20.4, *Bacillus amyloliquefaciens* 7.6, *Pseudomonas turukhanskensis* 14.7, *Microbacterium aerolatum* 14.4*, Agrobacterium fabacearum* 6.8*, Microbacterium neimengense* 10.2*, Enterobacter cancerogenus* 14.3*, Epilithonimonas hungarica* 19.2*, Shinella oryzae* 21.1*, and Pseudacidovorax intermedius* 2.1 showed higher nitrogenase activity than the reference diazotrophs *K. variicola* A3 and *A. brasilense* FP2 (**Figure 4D**). These findings validate that sorghum mucilage hosts a variety of diazotrophs capable of potentially aiding in nitrogen fixation[42].

### Mucilage enables efficient biological nitrogen fixation in sorghum

In Sierra Mixe maize landraces, the mucilage fosters a nitrogen-acquiring habitat for diazotrophs with low oxygen levels and an abundance of carbohydrates [13]. Using ARA, we tested whether sorghum mucilage could serve as a proper medium for nitrogen fixation. A panel of nine diazotrophs isolated from sorghum and maize mucilage along with other isolates from cereals (e.g., maize, rice (*Oryza sativa* L.), wheat (*Triticum aestivum* L.), and sugarcane) were used for this experiment (**Supplementary Table 8**). As a negative control, we included *A. brasiliense* FP10, a strain lacking nitrogenase activity yet exhibiting a growth rate equivalent to the wild-type FP2 strain [41]. To limit the abundance of environmental diazotrophic bacteria in the sorghum mucilage, we collected mucilage from sorghum plants grown in the greenhouse. We selected sorghum accession IS23992 because it produced numerous nodes with aerial roots in greenhouse and field conditions (**Figure 1 and Supplementary Figure 2**). ARA indicated significant nitrogenase activity in mucilage inoculated with the strains *Azotobacter vinelandii* DJ, a reference diazotroph for decades, and *Klebsiella variicola* A3 that we isolated sorghum mucilage (**Figure 5A**). However, all seven other strains tested exhibited limited nitrogenase activity, with only a *Klebsiella michiganensis* strain isolated from maize mucilage showing modest ARA activity. This experiment indicated that the sorghum aerial root mucilage provides a proper environment for some diazotrophs to fix nitrogen, particularly those isolated from mucilage.

**Fig 5.**
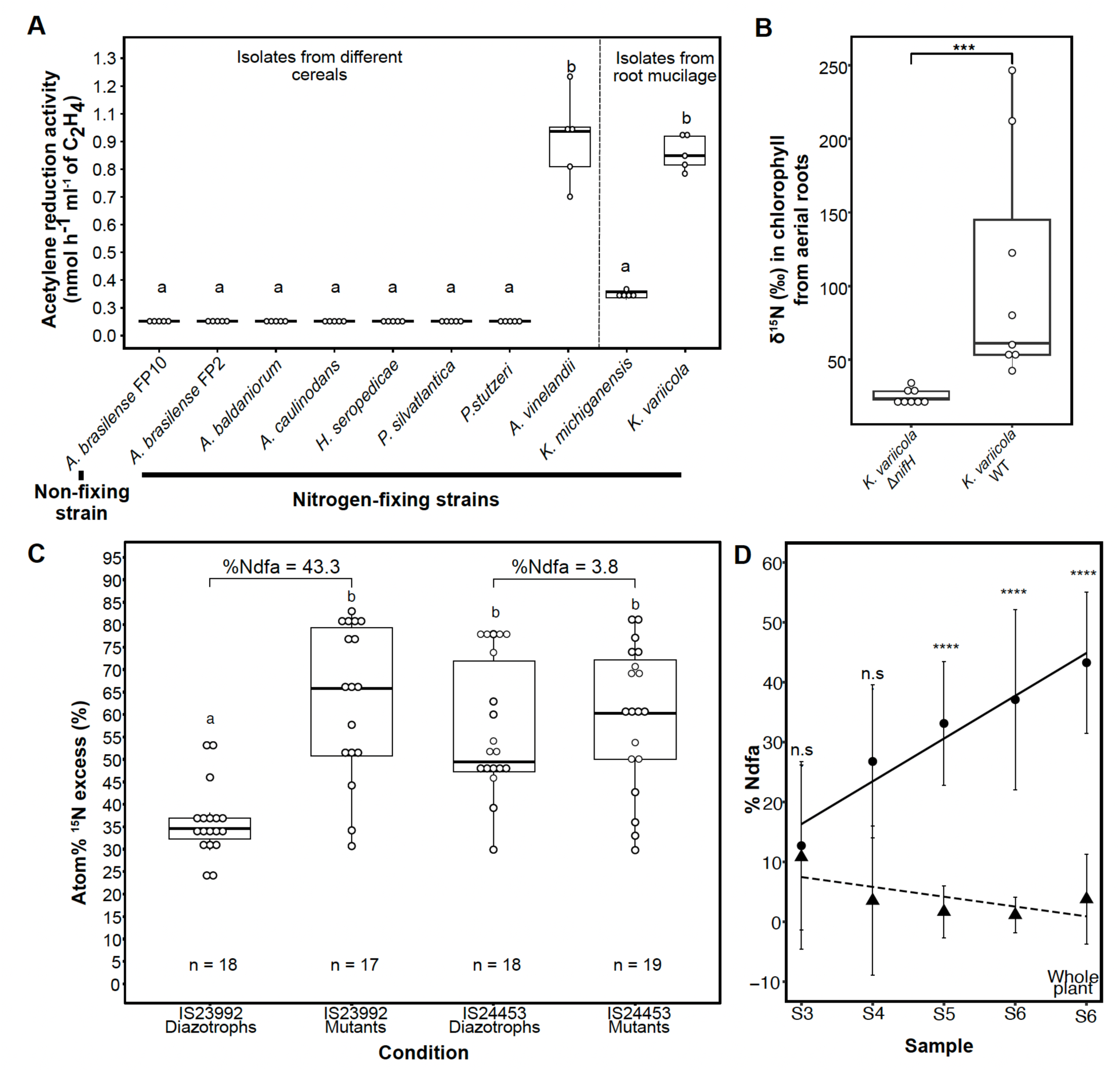
Biological nitrogen fixation on aerial roots from sorghum. A) Acetylene reduction assay in mucilage generated in the greenhouse. Mucilage was used as media to grow 9 nitrogen-fixing strains (*Azospirillum brasilense* FP2, *Azospirillum baldaniorum*, *Azorhizobium caulinodans*, *Azotobacter vinelandii* DJ, *Herbaspirillum seropedicae*, *Klebsiella michiganensis*, *Klebsiella variicola* A3, *Paraburkholderia silvatlantica*, and *Pseudomonas stutzeri*) and *Azospirillum brasilense* FP10 as a non-fixing control. Tukey’s honestly significant difference (HSD) test was performed in R, and significant group means are displayed as lower case letters (n = 6). B) Analysis of ^15^N gas feeding experiment on aerial roots of sorghum accession IS23992. Aerial roots were inoculated with *K. variicola* Δ*nifH* (negative control) or *K. variicola* A3 (positive control). *** indicates p-value ≤ 0.001 based on Wilcoxon signed-rank test (n = 8). C) ^15^N isotope dilution experiment on aerial roots of sorghum accessions IS23992 and IS24453. Accessions IS23992 and IS24453 were inoculated with the nine strains from the ARA experiment (diazotrophs), and three non-fixing mutant strains *Azospirillum brasilense* FP10, *Azotobacter vinelandii* DJ100 Δ*nifH*, *Azorhizobium caulinodans* ORS571 Δ*nifA*. Whole plants at the flowering stage were ground, and the average of three replicates was plotted. Percent Ndfa was calculated for both accessions using the plants inoculated with diazotrophs and using as reference the plants inoculated with mutant strains. An ANOVA was performed to compare means between treatments and genotypes using the package multcompView. D) Scatterplot of the percentage of nitrogen derived from the atmosphere (%Ndfa) in different sorghum samples from the accessions IS23992 and IS24453, indicated with circles and triangles, respectively. Solid and dotted lines are regression lines for IS23992 and IS24453, respectively. Samples were collected every two weeks, starting at stage 3 (Samples S3 to S5) and after flowering (S6). Error bars display standard deviations. Pair-wise comparisons were made with Wilcoxon’s test;**** means p-value ≤ 0.0001 and n.s. Indicates non-significant.

To test the plant’s ability to assimilate nitrogen fixed by diazotrophs, we conducted a ^15^N_2_ gas feeding experiment. Aerial roots producing mucilage were collected from the greenhouse-grown sorghum plants of the accession IS23992. Based on the ARA results, we inoculated the roots with wild-type fixing *Klebsiella variicola* A3 or its non-fixing Δ*nifH* mutant strain as a negative control[42]. Isotope-ratio mass spectrometry (IRMS) was used to quantify the δ^15^N in chlorophyll from sorghum aerial roots. Roots inoculated with wild-type fixing *K. variicola* exhibited significant incorporation of ^15^N in plant chlorophyll compared to aerial roots inoculated with the non-fixing strain (**Figure 5B**). To provide additional evidence of fixed nitrogen uptake, we conducted a ^15^N isotope dilution assay. Two sorghum accessions, IS23992 and IS24453, exhibiting high and low numbers of aerial roots, respectively, were grown in pots in the greenhouse. These accessions were inoculated either with a panel of diazotrophs from the ARA experiment or three non-fixing mutants (**Supplementary Table 8**). Atom%^15^Nexcess was evaluated by IRMS in leaves collected at four vegetative stages (S3, S4, S5, and S6) and whole plant samples at S6. Significantly lower Atom%^15^Nexcess values were observed in the IS23992 accession, which produces many aerial roots, when inoculated with nitrogen-fixing bacteria compared to plants inoculated with the non-fixing controls. Atom%^15^Nexcess values of the IS24453 accession, which makes few aerial roots, inoculated with fixers or non-fixers, were similar to those of the IS23992 genotype inoculated with non-fixers (**Figure 5C**). Based on these ^15^N dilution data, we evaluated the percentage of nitrogen derived from the atmosphere (%Ndfa) at these four stages of sorghum development (**Supplementary Figure 5**). %Ndfa increased steadily over time, reaching 43% on whole plants at S6 (**Figure 5D**). Even though all the techniques to quantify nitrogen fixation (ARA, ^15^N_2_ gas feeding, and ^15^N isotope dilution) were used in the greenhouse and have their own biases, our results collectively indicate that sorghum accessions with a large number of aerial roots can support an active community of diazotrophs and obtain a significant fraction of their nitrogen from the biological nitrogen fixation.

## Discussion

### Some sorghum accessions produce abundant and thick aerial roots under humid conditions

Previous studies have reported variations in the number of nodes adventitious nodal roots among different sorghum genotypes. For example, the Chinese landrace Sansui produces 6-8 nodes with aerial roots, while the elite cultivar Jiliand only has a single node with roots [43]. Our study based on select sorghum accessions from different sources such as the sorghum mincore panel, the sorghum association panel and ICRISAT confirmed significant variation in the number of nodes with aerial roots (**Figure 1A**). We observed differences in diameter and mucilage production in these accessions, and, as in maize, a positive correlation among these traits was confirmed [13]. The presence of aerial roots in other members of the Andropogoneae and Paniceae tribes, such as sugarcane and foxtail millet, suggests a common evolutionary origin of this trait [18]. It will be interesting to explore if similar genes control this trait across these species.

Water availability, hypoxia, and humidity impact plant growth and development by producing phytohormones, including abscisic acid and ethylene [44]. In the case of rice, high humidity stimulates the development of lateral roots, which was attributed to increased levels of ethylene [45–47]. Abscisic acid also controls plant responses to water availability [48]. Under high humidity, we observed more nodes with aerial roots across sorghum genotypes, possibly by regulating phytohormones like ethylene and abscisic acid. The accession IS24453 failed to develop aerial roots in the initial experiment, where humidity remained at base levels. However, it eventually developed aerial roots upon increasing humidity, likely due to increased phytohormone levels (**Figure 2**). Subsequent experiments should prioritize quantifying these components to mitigate unidentified experimental variables that may influence aerial root development.

### Some sorghum accessions produce abundant mucilage from aerial roots

Plants secrete mucilage from various tissues to adapt to abiotic and biotic stresses. In underground roots, mucilage production is carried out by border cells. In the case of the maize landraces, the mucilage from their aerial roots resembles the mucilage commonly found at the tips of underground roots. Pankievicz *et al*. [15] showed the abundance of border cells in the mucilage from the aerial roots of Sierra Mixe maize. They noted differences in border cell sizes compared to border cells from underground roots. This result explained the difference in mucilage production between the two types of roots [15]. Recently, a study involving 146 sorghum accessions described the generation of mucilage from the aerial roots[49]. Our data corroborated these sorghum findings, showing differences between border cell morphologies between underground and aerial roots and mucilage production (**Supplementary Figure 3 and Figure 3A**). Based on these findings, we hypothesize that the border cells in sorghum and maize exhibit greater similarity than those in underground roots. The difference in quantity and size explains the volume generated by each type of root. Additionally, we hypothesize these specialized cells are more abundant and functionally active in younger roots, as observed in maize aerial roots.

Carbohydrates are the primary components of both underground and aerial root mucilage [13,14,50,51]. In this study, we investigated the carbohydrate composition of sorghum mucilage and showed the presence of similar monosaccharides in maize and sorghum aerial root mucilage, albeit in different proportions (**Figure 3**). These findings are consistent with a prior study on maize genotypes, which revealed changes in the carbohydrate profile based on their geographic locations and agroecological systems [51]. Wolf *et al.* also analyzed the monosaccharide composition in mucilage from sorghum aerial roots, and we found that both studies reported a similar range of values [28]. To fully understand variations in monosaccharide composition in sorghum mucilage, a broader range of environments and genotypes needs to be considered. Interestingly, our analysis did not identify the presence of fructose in the sorghum mucilage, as reported in other studies (**Figure 3B**) [49]. This mucilage composition may suggest that environmental factors significantly influence maize and sorghum mucilage. However, the genetic aspect cannot be disregarded entirely, as evidenced by our principal component analysis, which displayed a clear separation based on species (**Figure 3**). Overall, inter-species differences are more pronounced than inter-accession differences (**Figure 3B and 3D)**, supporting reports that highlight variations in the microbiomes of cereals [52]. Additionally, there must be differences within genotypes, as observed in maize underground mucilage[51].

### Sorghum aerial roots mucilage hosts specific microbial communities containing diazotrophs

Several studies have documented variations in plant microbiome composition based on host and environmental factors [53–56]. Our microbiome survey revealed that location and host influence microbial composition in the aerial root mucilage (**Figure 4**). Certain plant enzymes that break down mucilage into simple sugars are expressed in the Sierra Mixe landraces and may play a role in selecting specific microbes. The microbiome may contribute to mucilage degradation through specialized enzymes [15]. Pseudomonadota, Acidobacteriota, Cyanobacteriota, and Bacillota were the most abundant phyla in sorghum, aligning with findings from other sorghum microbiome studies [57]. Our study sheds light on the microbial profiles of sorghum and maize for nitrogen fixation and offers valuable strains for engineering diazotrophs and enhancing nitrogen fixation. These resources will serve as a foundation for investigating microbial community interactions in the mucilage. Such insights could pave the way for inoculation strategies with diazotrophic consortia and/or engineered diazotrophs.

Sorghum growth promotion enabled by inoculation with diazotrophs has been reported several years ago [58–60]. There was evidence that sorghum could acquire variable amounts of nitrogen from the air after inoculation with diazotrophs or in association with native bacterial communities [61,62]. However, these reports provided no information about the ability of these genotypes to produce aerial roots or mucilage. Our study provides evidence that the production of aerial roots and mucilage is a mechanism in sorghum that facilitates association with diazotrophs, enabling the plant to obtain nitrogen from the atmosphere through biological nitrogen fixation. Microbial communities can thrive in plant mucilages due to the abundance of carbon, and diazotrophs often benefit from the microaerobic conditions. The mucilage viscosity is strikingly similar to nitrogen-free semi-solid media, which is classically used to isolate diazotrophs from non-legumes [63]. The presence of diazotrophs in sorghum aerial root mucilage was confirmed in our study using bacterial isolation in nitrogen-free semisolid media and nitrogenase activity assays.

### Efficient nitrogen fixation occurs in sorghum aerial root mucilage

Biological nitrogen fixation in cereals can reduce agriculture’s dependence on synthetic nitrogen-based fertilizers. Our research has shown that mucilage is crucial in creating a favorable environment for diazotrophs to thrive, enabling plants to acquire nitrogen (**Figure 5A, B, and C**). Two independent studies have revealed that specific diazotrophs form associations with the underground roots of sorghum plants, delivering nitrogen to the plant. We observed this unique specificity as only the diazotrophs isolated from mucilage exhibited higher nitrogenase activity than those isolated from other grasses, considered the benchmark for studying nitrogen fixation in cereals (**Figure 5A**). The biological nitrogen fixation in sorghum was previously studied through ARA, ^15^N isotope dilution, and ^15^N abundance assays [62,64,65]. This study used three complementary approaches (ARA, ^15^N_2_ gas feeding, and ^15^N isotope dilution) to provide evidence of efficient nitrogen fixation in sorghum aerial roots (**Figure 5**). While these methods can suffer from inherent biases, they collectively point towards efficient nitrogen fixation in the sorghum mucilage and the acquisition of this nitrogen by the plant. The mucilage offers a favorable environment for diazotrophs; therefore, obtaining mucilage or mucilage-producing plants devoid of diazotrophs can be challenging. Thus, for these experiments, rather than using non-inoculated mucilage or plants as negative controls, we favored inoculation with non-fixing mutants to exclude as much as possible diazotrophs from the environment. ARA revealed that diazotrophs can actively fix nitrogen in this environment (**Figure 5A**). However, ARA cannot prove that the plant can acquire this nitrogen. This was demonstrated in ^15^N_2_ gas feeding and ^15^N isotope dilution (**Figures 5B and 5C**). Previous field studies using ^15^N isotope dilution in sorghum suggested that diazotrophs can provide between 36 and 78 % of nitrogen depending on the edaphoclimatic conditions [62]. In this study using ^15^N isotope dilution in greenhouse conditions, we observed that sorghum with aerial roots can acquire up to 43% of the atmospheric nitrogen (**Figure 5C**). ^15^N dilution can be biased due to the segregation of ^15^N within the soil, and different plants may acquire nitrogen from different soil layers. In our case, the sorghum root system filled the pots, acquiring nitrogen from all the soil layers and alleviating this concern. However, we should be careful to extrapolate results from greenhouse experiments to field conditions where the environment will undoubtedly positively or negatively affect this plant-microbe relationship.

### Sorghum,maize and their relatives with abundant and thick aerial roots as promising avenues to reduce the use of synthetic nitrogen fertilizers

Farmers in Oaxaca, Mexico, have preserved and cultivated the Sierra Mixe maize landraces for centuries. Field experiments indicated that soils with more than ten years of maize cultivation showed higher %Ndfa values than those with zero to one year of maize history. This consistent cultivation has promoted biological nitrogen fixation in these maize landraces[13]. Consequently, this selection process has significantly enhanced biological nitrogen fixation. To our knowledge, the sorghum accessions we identified have undergone minimal selection for biological nitrogen fixation, which may explain the differences in %Ndfa between sorghum and maize. By adopting modern cultivation practices, we may achieve increased levels of %Ndfa in sorghum.

Sorghum is renowned for its resilience to drought, which might restrict the areas suitable for exploiting biological nitrogen fixation from aerial roots with mucilage. In maize, drought-adapted cultivars can release mucilage that retains water effectively [66]. Therefore, sorghum accessions that produce aerial roots and that originate from drought-prone regions may also produce mucilage under drought conditions. However, (sub)tropical regions or areas with high relative humidity might be ideal for these sorghum accessions.

This study examined the diversity within the minicore and ICRISAT sorghum collections and identified accessions that developed aerial roots. Sorghum, maize, and sugarcane are all members of the Andropogoneae tribe, and genera from the Paniceae tribe (Setaria and Pennisetum) are known to develop brace roots [18]. In sugarcane, thin roots emerging from nodes have been observed [67]. During a screening of Setaria, we discovered *Setaria italica* and *Setaria magna* accessions with thin aerial roots lacking mucilage. Additionally, we found that Johnsongrass (*Sorghum halepense*) and sudangrass (*Sorghum bicolor* × *Sorghum arundinaceum*) developed aerial roots with mucilage (**Supplementary Figure 6**). Notably, these grasses exhibited thicker roots and mucilage. Our research indicates thicker roots are necessary for mucilage production, as seen in maize and sorghum. Two loci associated with aerial root diameter in sorghum were reported by Wolf et al. [39]. Based on the observations of aerial root formation in other grasses, additional screenings of germplasm collections can provide more information on the genetic basis of the aerial root formation and aerial root diameter.

To achieve efficient nitrogen fixation in cereals, synthetic biology offers exciting approaches through transferring nitrogenase genes to plants, engineering root nodule symbiosis, or remodeling associative diazotrophs [68,69]. While exploring these approaches is undeniably valuable, it is important to explore the untapped potential present in the natural diversity of plants to identify and breed superior plant hosts for diazotrophs using non-transgenic methods. Certain maize landraces in Sierra Mixe can obtain significant amounts of nitrogen from the atmosphere through their aerial roots. Our research suggests that a similar trait is present in sorghum, and investigating the natural diversity in cereal crops for nitrogen fixation provides a near-term, non-transgenic pathway to enhance biological nitrogen fixation in these crops. Leveraging our genetic understanding of this trait, we can engage in breeding efforts in sorghum and maize to promote aerial development and mucilage secretion [28]. These natural traits are quite promising to reduce the dependence on synthetic nitrogen fertilizers for cereal crops and improve the sustainability of our agriculture for food, feed, fodder and biofuel production.

## Materials and methods

### Plant material and phenotyping

Sorghum accessions were selected from the sorghum minicore collection [25] and augmented with accessions kindly provided by Vetriventhan Mani and Vania C. R. Azevedo at the International Crops Research Institute for the Semi-Arid Tropics (ICRISAT, Hyderabad, Telangana, India) (**Supplementary Table 1**). The plants were cultivated in C1200 nursery pots containing Metro-Mix® 360 SUN-COIR™ (Burton, Ohio) medium in the Walnut Street Greenhouses at the University of Wisconsin-Madison with temperatures at 25 °C during the day and 17 °C at night. The plants were started on a long daylight regimen (14/10), and once they were 1-1.5 m tall, they were switched to short days (12/12) due to their photoperiod sensitivity. Relative humidity was set at 20%. Sorghum plants were watered with municipal water or NPK fertilizer (20-10-20). At the flowering stage, when aerial roots were visible on many nodes, the root diameter of three roots at the top node was measured with a caliper, as well as the number of nodes forming aerial roots and the number of aerial roots formed on the top node. Overhead water was applied, and 30 minutes after the water application, the mucilage volume was quantified from three young and three old aerial roots with similar lengths, of approximately 10 cm. Mucilage was collected into 2 ml tubes by simply removing it from the base to the tip of the root. The same accessions were grown at the University of Wisconsin-Madison West Madison Agricultural Research Station (43.0610329663, -89.5325872441) and the University of Florida North Florida Research and Education Center - Suwannee Valley near Live Oak, FL (30.313277, -82.902158) in 1.5 m rows, with 10 cm between plants, and 76 cm between rows. Only the number of nodes and root diameter were phenotyped in the field experiments.

### Greenhouse experiments on the influence of humidity levels

Sorghum accessions IS23992 and IS24453 were grown in individual 6-gallon (23-L) pots (Custom-Tainer 2800C-14") with Pro-Mix LP15 medium (Quakertown, Pennsylvania) at the Walnut Street Greenhouses at the University of Wisconsin-Madison. Greenhouse temperature was set at 28 °C during the day and 25 °C at night. Plants were grown under a light regime of 12-hour photoperiod from 6 am to 6 pm. Greenhouses were kept at high (75%) or low relative humidity 30%, controlled with a humidifier system (Smart Fog, Reno, Nevada). The phenotypes evaluated included number of nodes with roots, the number of roots at the top node, and the root diameter. The measurements were obtained with the previously described procedure.

### Bacterial growth conditions

Twelve strains were used for different experiments (**Supplementary Table 8**). *Azospirillum brasilense* FP2 and FP10 (*nifA^-^*) were kindly provided by Fábio Pedrosa and Emanuel M. de Souza from UFPR-Brazil[41]. The *Klebsiella variicola* Δ*nifH* mutant was developed by Maya Venkataraman (UW-Madison). *Azospirillum baldaniorum* Sp 245^T^, *Klebsiella variicola* A3, *Klebsiella michiganensis* A2, *Azorhizobium caulinodans* ORS571^T^, *Stutzerimonas stutzeri* A1501, *Paraburkholderia silvatlantica* SRMrh-20^T^, *Azotobacter vinelandii* DJ, *Herbaspirillum seropedicae* SmR1, *Azorhizobium caulinodans* ORS571 Δ*nifA* were grown in LB medium, except for *A. vinelandii* DJ and *A. vinelandii* DJ100 Δ*nifH*, which were grown in Burk’s sucrose medium[70,71]. The cells were grown overnight at 30 °C while shaking at 130 rpm.

### Mucilage sugar composition

Root mucilage from accessions IS11026, was collected from cut aerial roots of different sorghum genotypes from the greenhouse experiment. The roots underwent a 24-hour water exposure in a 50 ml tube, and the mucilage from two separate experiments was sent to the Complex Carbohydrate Center Research Center (CCRC) at the University of Georgia, Athens, USA. Glycosyl composition analysis was performed by combined gas chromatography/mass spectrometry (GC/MS) of the per-*O*-trimethylsilyl (TMS) derivatives from the monosaccharide methyl glycosides produced by acidic methanolysis as described previously [13]. The analysis included rabinose, rhamnose, ribose, fucose, xylose, glucuronic acid, galacturonic acid, mannose, galactose, glucose, *N*-acetyl mannosamine, *N*-acetyl galactosamine, *N*-acetyl glucosamine were measured but only monosaccharides that were detected are reported.

### Total free sugar quantification

Mucilage was collected at two time points from the sorghum accession IS23992 grown under greenhouse conditions. Benedict’s reaction was used to quantify free sugars, as described in Hernandez-Lopez *et al*.[27]. Benedict’s reagent (700 μl) was incubated with sorghum mucilage (30 μl) for 10 min at 95 °C. Subsequently, the sample was centrifuged at 10,000 *g* for 5 min. The supernatant was collected, and the absorbance at 740 nm was determined using a BioRad SmartSpect^TM^ Plus spectrophotometer. Sugars were quantified using a calibration based on solutions with different concentrations of D-glucose.

### Metabolomic analyses

Metabolome analysis of mucilage samples from sorghum accessions IS29091 and IS2241, as well as maize landraces CB017456 and GRIN 19897, was conducted at the West Coast Metabolomics Center at UC Davis following their standard protocols (https://metabolomics.ucdavis.edu/). A minumum of 75 mg of mucilage was sent per sample (Supplementary Table 3). A primary metabole analysis was performed to detect sugar phosphates, amino acids, hydroxyl acids, free fatty acids, purines, pyrimidines, aromatics compounds, and unidentified compounds. Metabolites were quantified by peak heights, normalized, and processed with MetaboAnalyst[72] (https://www.metaboanalyst.ca/). This algorithm offers several different methods. Sparse partial least squares - discriminant analysis and significance analysis of microarray were used to analyze the data.

### Microscopy

Border cells from underground roots were collected from the root tip of six-day-old seedlings from sorghum accession IS23992 that grew in 1.2% agar at 24 °C in a growth chamber. Border cells from aerial roots were collected from mucilage produced from the humidity study. The mucilage was triggered by spraying the aerial roots directly with municipal water 5 times every 5 minutes. The mucilage was collected after 1 hour after the final spraying. Approximately 5 μl of mucilage were spread on a slide and combined with 5 μl of distilled water, 10 μl of SYTO green (5 μM), and 10 μl of propidium iodine (10 μg/mL). The cell suspension was observed under a Leica DMi8 model S inverted fluorescence microscope. GFP and mCherry filters were used to examine SYTO green and propidium iodide fluorescent signals, respectively. Light microscopy was used to obtain images of border cells from both underground and aerial roots. ImageJ - Fiji (version 2.9.0) [73] was used to measure the area and length of the border cells.

### Microbiome study and 16S rRNA amplicon sequencing

DNA was isolated from the mucilage samples from sorghum and maize using the FastDNA™ SPIN Kit for Soil (MP Biomedicals, Irvine, California) following the manufacturer’s instructions. DNA was quantified using a Nanodrop One (Thermofisher Scientific, Waltham, Massachusetts). The V3-V4 region of the *16S rRNA* gene was amplified with the standard bacterial primers 335F and 769R [74] with Go*Taq* Mastermix (Promega, Madison, Wisconsin) according to the manufacturer’s instructions. Amplification reaction volumes were 50 µl containing 10 pmol of each primer and 4 ng of template DNA. Thermocycler conditions were as follows: 95°C for 1 minute, then 35 cycles of 95°C for 10 s, 55°C for 30 s, and 72°C for 30 s, with a final step at 72°C for 5 min. PCR products were separated on an agarose gel and extracted using the GeneJET Gel Extraction Kit (Thermofisher Scientific). Novogene (Santa Clara, California, USA) conducted library preparation and sequencing. Amplicons were sequenced with an Illumina paired-end platform generating 250 bp paired-end raw reads. OTUs abundance was normalized using a standard sequence number corresponding to the sample with the fewest sequences. Sequences exhibiting a similarity of ≥97% were grouped into identical OTUs. A representative sequence was chosen for each OTU to undergo subsequent annotation. Principal component and beta diversity analyses were performed based on these normalized data. These data were processed with QIIME2 (Version 1.7.0) [75] and displayed with R software (Version 2.15.3) package ggplot2 (Version 3.4.3.) [76]. Sequencing data are available at the Short Read Archive of NCBI under BioProject: PRJNA1009066.

### Isolation of bacteria from sorghum samples

Mucilage samples were collected from a West Madison Agricultural Research Station field trial in 2022 from mucilage after rain (**Supplementary Tables 5 and 7**). The sampling was conducted after a continuous rainy weekend with an accumulated rainfall of around 60 mm. In the field, we looked for homogeneous mucilage production in the sorghum association panel [28,39]. A sample was composed of a mix of two replicates from the panel. Mucilage (2-3 mL per plant) was collected with a 1-mL pipette in a 15 mL tube. The samples were kept cold (0 °C) until analysis. The mucilage samples were added to BMGM semisolid medium and incubated at room temperature for one week [77]. After that, the microaerophilic pellicles that formed were transferred to a fresh medium and incubated for the same period seven more times. After the final incubation period, solid BMGM medium supplemented with 0.5 g/l of yeast extract was inoculated with the bacteria. A total of 164 isolates were screened by clonal strains using the Box-PCR approach with the A1 primer (5′-CTACGGCAAGGCGACGCTGACG-3′) and subjected to agarose gel electrophoresis (1.2%) [78]. Identical profiles were excluded, and 103 non-clonal strains were kept. The bacterial strains were identified by sequencing 16S rRNA genes using the primers 27F (5’- AGAGTTTGATCCTGGCTCAG-3’) and 1492R (5’-GGTTACCTTGTTACGACTT-3’)[79]. The PCR was initiated with a denaturation cycle at 95°C for 10 minutes, followed by 35 cycles consisting of denaturation at 95°C for 30 seconds, annealing at 55°C for 30 seconds, and extension at 72°C for 1 minute and 30 seconds. The process concluded with a final extension at 72°C for 10 minutes. PCR product was sequenced in an Applied Biosystems 3500 genetic analyzer at the Functional Biosciences facilities. Sequence Scanner v.2.0 (Informer Technologies, Inc.) was used for quality assessment, and the contigs of almost complete 16S rRNA sequences were assembled in BioEdit v.7.7[80]. The sequences were compared against those deposited within the 16S rRNA reference database of GeneBank using the BLASTn tool [81]. The maximum likelihood phylogenetic tree was constructed using MEGA 11 software [82]. The aligning used 103 different 16S rRNA gene sequences from the bacteria isolated from the sorghum mucilage.

### Acetylene reduction assays (ARA) to quantify nitrogenase activity

Nitrogenase activity was evaluated in ten bacterial strains (**Supplementary Table 8**) grown in sorghum mucilage collected from the greenhouse. The bacteria were grown in 3 ml of LB medium overnight at 30 °C and shaken at 180 rpm. The culture was centrifuged at 5000 *g* for 5 minutes, and the pellet was resuspended in 1 ml of sterile water. Then 30 µl (OD_600nm_ = 0.1) were inoculated in sterile vials containing 3 ml of mucilage. Vials were capped and incubated for 24 hours at 30 °C. The cap was replaced with a crimp cap, and 1 ml of air was replaced with acetylene, followed by a 3-day incubation period at 30 °C. Gas Chromatography - Flame Ionization Detection was performed in a GC-2010 (Shimadzu) with an HS20 autosampler by injecting 1 ml of air sample. The injector and the detector were set at 200 °C with a flow rate of 86.6 ml/min nitrogen carrier gas. The temperature of the column was 100 °C. Controls included a vial with acetylene to check for traces of ethylene and *A. brasilense* FP2 without fixing activity. The indirect measure of the nitrogenase activity was expressed in nmol of ethylene produced per hour per mL of mucilage. ARA in the 34 isolated diazotrophs was performed similarly but using a semi-solid BMGM medium.

### 15N2 gas feeding experiments

Aerial roots from sorghum accession IS23992 were collected in 150 ml sterile flasks. These roots were obtained from greenhouse-grown plants with copious amounts of mucilage. *K. variicola A3* and the *K. variicola* Δ*nifH* mutant were grown overnight at 30 °C and shaken at 180 rpm in LB medium. The culture underwent centrifugation at 5000 *g* for 5 minutes, and the resulting pellet was then resuspended in 1 ml of sterile water. Bacteria were directly inoculated onto the surface of mucilage-coated roots (100 µl, OD_600nm_ = 0.1). Flasks were injected with ^15^N_2_ gas (98%) to reach a concentration of 15% (v/v), and flasks were incubated for three days at room temperature (20°C). Non-fixing bacteria were included as negative controls to confirm the absence of other sources of ^15^N in the system. The roots with mucilage were excised from the plant for practicality and proper incubation in ^15^N_2_ gas. To determine the ^15^N enrichment specifically in plant tissue, chlorophyll was extracted by peeling the green cell layers from the aerial roots after removal of mucilage. The extraction followed the protocol described by Kahn *et al.* [83]. The samples were analyzed by isotope ratio mass spectrometry (IRMS) at the Department of Soil Science at the University of Wisconsin-Madison to evaluate the δ^15^N of the samples.

### 15N isotope dilution experiments

A total of 40 sorghum plants from each of the two accessions, IS23992 and IS24453, were grown in 6-gallon (23-L) pots at the Walnut Street Greenhouses in two adjacent greenhouse rooms to prevent contamination between the two types of bacterial inoculants (fixing and non-fixing). An equal number of plants were placed in the two treatments, and standard humidity conditions were set (relative humidity 50%) with a humidifier system (Smart Fog, Reno, Nevada). After four weeks post-germination, pots were fertilized weekly with 500 ml of Hoagland’s No.2 Basal Salt mixture (1.34 g per litter) and 80% ammonium-^15^N sulfate (^15^NH_4_)_2_SO_4_ as the only source of nitrogen. Five weeks after germination, aerial roots of plants in one room were inoculated with a mixture of nine fixing strains (OD_600_ = 0.1), and the aerial roots on the plants in the other room was inoculated with three non-fixing strains (OD_600_ = 0.1) (**Supplementary Table 8**). Plants were sprayed with 250 ml of the inoculum resuspended in a phosphate-buffered saline (PBS) buffer using a backpack sprayer. The third leaf counting from the top was collected before inoculation (stage 3) and then three times every two weeks (stages 4 and 5). At flowering (stage 6), whole plants were collected, ground, and homogenized to a fine powder. Quantification of the samples’ ^15^N content above the atmospheric concentration of N_2_ (Atom% ^15^N excess) was measured by isotope ratio mass spectrometry (IRMS) at the Department of Soil Science at the University of Wisconsin-Madison. The values of whole-plant samples were averaged. The percentage of nitrogen derived from the atmosphere (%Ndfa) was calculated according to the equation described in van Deynze *et al*.[13]. The fixers were the accessions inoculated with diazotrophs, whereas the references were those inoculated with the mutant strains.

### *Setaria* growth conditions

*Setaria* accessions PI43384 and PI689086 were cultivated at the Walnut Street Greenhouses under high relative humidity (80%) using the same system as described for the humidity experiment with sorghum. The plants were grown in 1-gallon (3.8-L) Pro Cal Premium Nursery Pots with Pro-Mix LP15 medium from Quakertown, Pennsylvania. Plants were maintained at 28 °C during the day and 25 °C at night, with a 12-hour photoperiod from 6 am to 6 pm. The phenotype evaluated included the number of nodes with roots.

### Raw data

All raw data are available at DOI: 10.6084/m9.figshare.25193696

### Statistical analyses

All statistical analyses were conducted in R version 4.2.1 and 2.15.3[84]. ANOVA and Tukey’s HSD were conducted with the package ggpubr (Version 0.6.0) and multcompView (Version 0.1-9). Wilcoxon’s test was also calculated with package ggpubr (Version 0.6.0). Microbiome studies were performed with QUIIME2 (Version 1.7.0) [85].

## Acknowledgments

We thank Sanhita Chakraborty and Kimberly Gibson for their valuable input and engaging discussions, Alyssa Davis and Julio Quiñones for their technical support, and Ben Broughton and Mike Boyette at the UF North Florida Research & Education Center-Suwannee Valley for managing the field plots. We thank Janet Hedtcke, the field manager at UW-Madison Agricultural Research Station. We thank Maya Venkataraman and Brian F. Pfleger for providing the *K. variicola* Δ*nifH* strain, David Brenner for providing us with seeds from different *Setaria* accessions, and Daniel Carmeli for providing pictures from Johnsongrass and sudangrass. Funding for this project was provided by the United States Department of Energy (DOE), Office of Science (BER), grant no. DE-SC0021052 (JMA, WV).

---

**Suppl. Fig 1.**
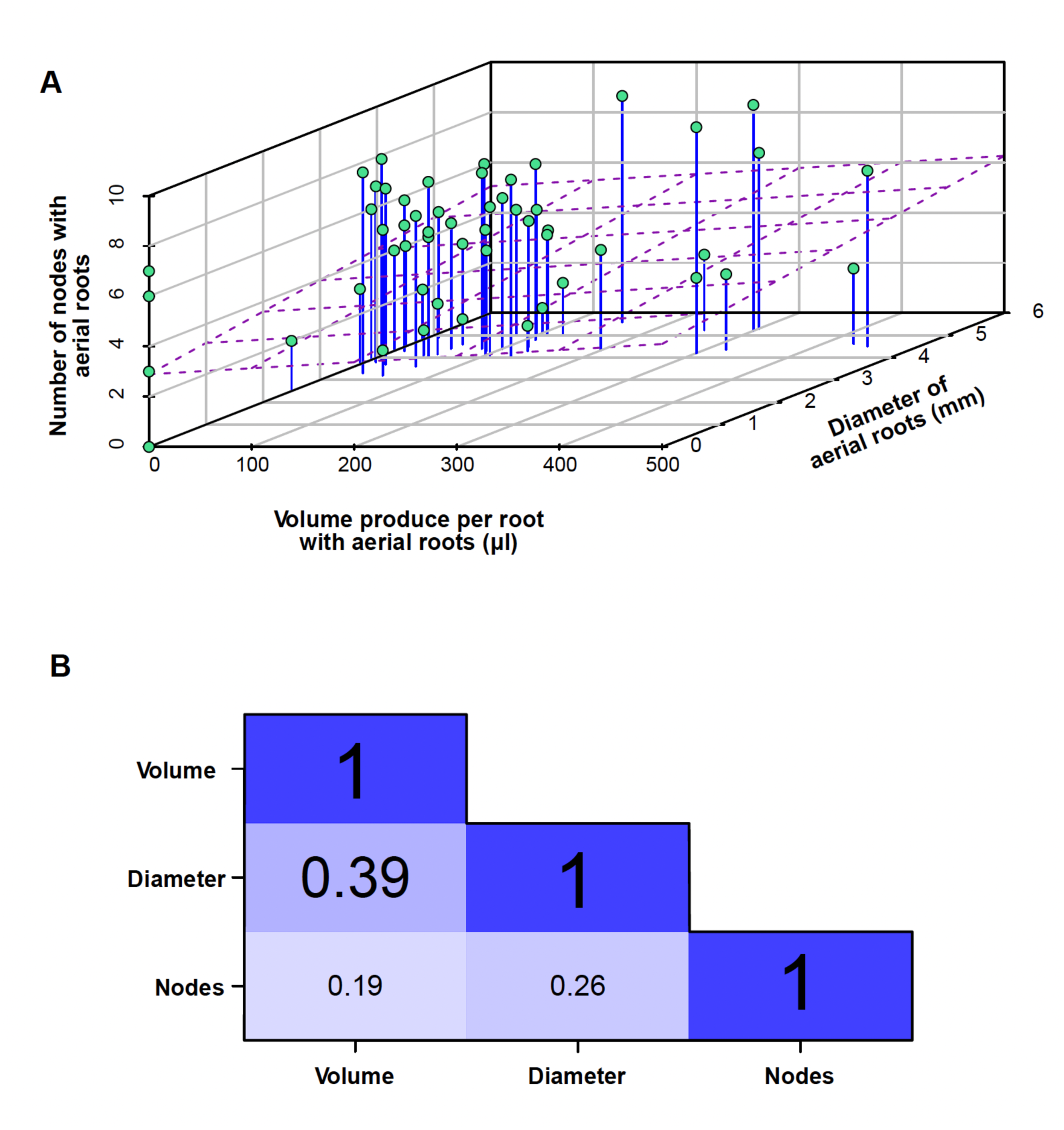
Trait correlations. A) Scatter plot shows the correlation between the number of nodes, mucilage volume, and diameter. A 3D regression line is dot plotted (purple) to observe the correlation among the traits. B) The correlation matrix for the number of nodes, mucilage volume, and diameter reveals a modest positive correlation across all traits. The correlation coefficient (r) values indicate the correlation strength between each trait. The correlation plot and values were calculated with the packages “scatterplot3d” [86] and “psych” [87], respectively.

**Suppl. Fig 2.**
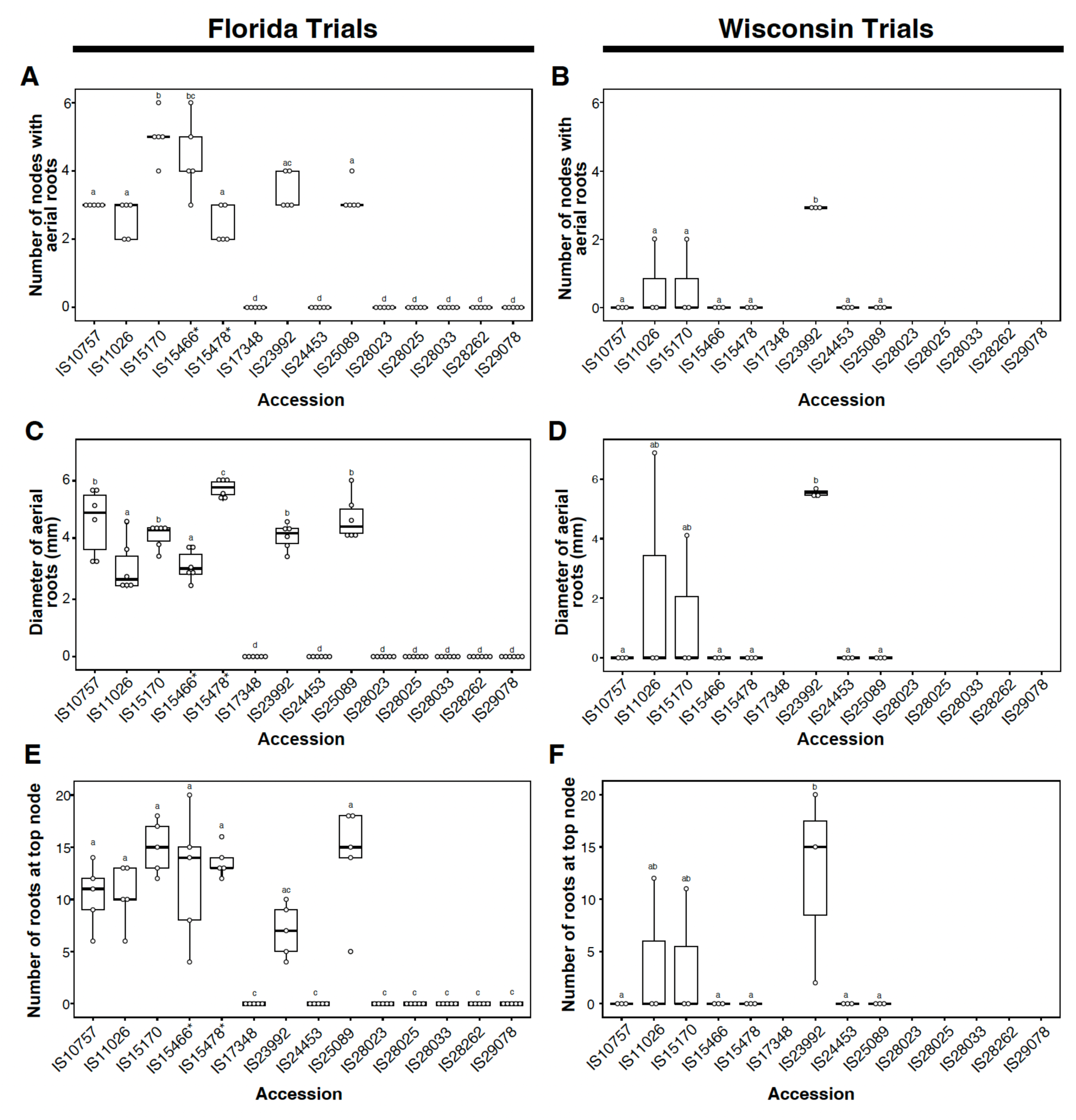
Sorghum screening of selected genotypes in two locations. A) Number of nodes in different sorghum accessions grown at the University of Florida during 2021 (n = 5). Generally, two groups can be observed, one with nodes with aerial roots and another without. B) Number of nodes in different sorghum accessions grown at the University of Wisconsin during 2022 (n = 3). Only specific accessions developed fewer aerial roots compared with the conditions in Florida. C) Diameter (mm) of aerial roots of different sorghum accessions grown at the University of Florida (n = 5). The accessions with aerial roots have thick diameters. D) Diameter (mm) of aerial roots in different sorghum accessions grown at the University of Wisconsin during 2022 (n = 3). Thickness was still observed in the accession with aerial roots. E) Number of aerial roots at the top node in different sorghum accessions grown at the University of Florida (n = 5). F) Number of aerial roots at the top node in different sorghum accessions grown at the University of Wisconsin during 2022 (n = 3). * Data from these accessions were obtained under a low nitrogen regimen. A subset of data for the Florida plots were obtained from the Wolf *et al.* study [28].

**Suppl. Fig 3.**
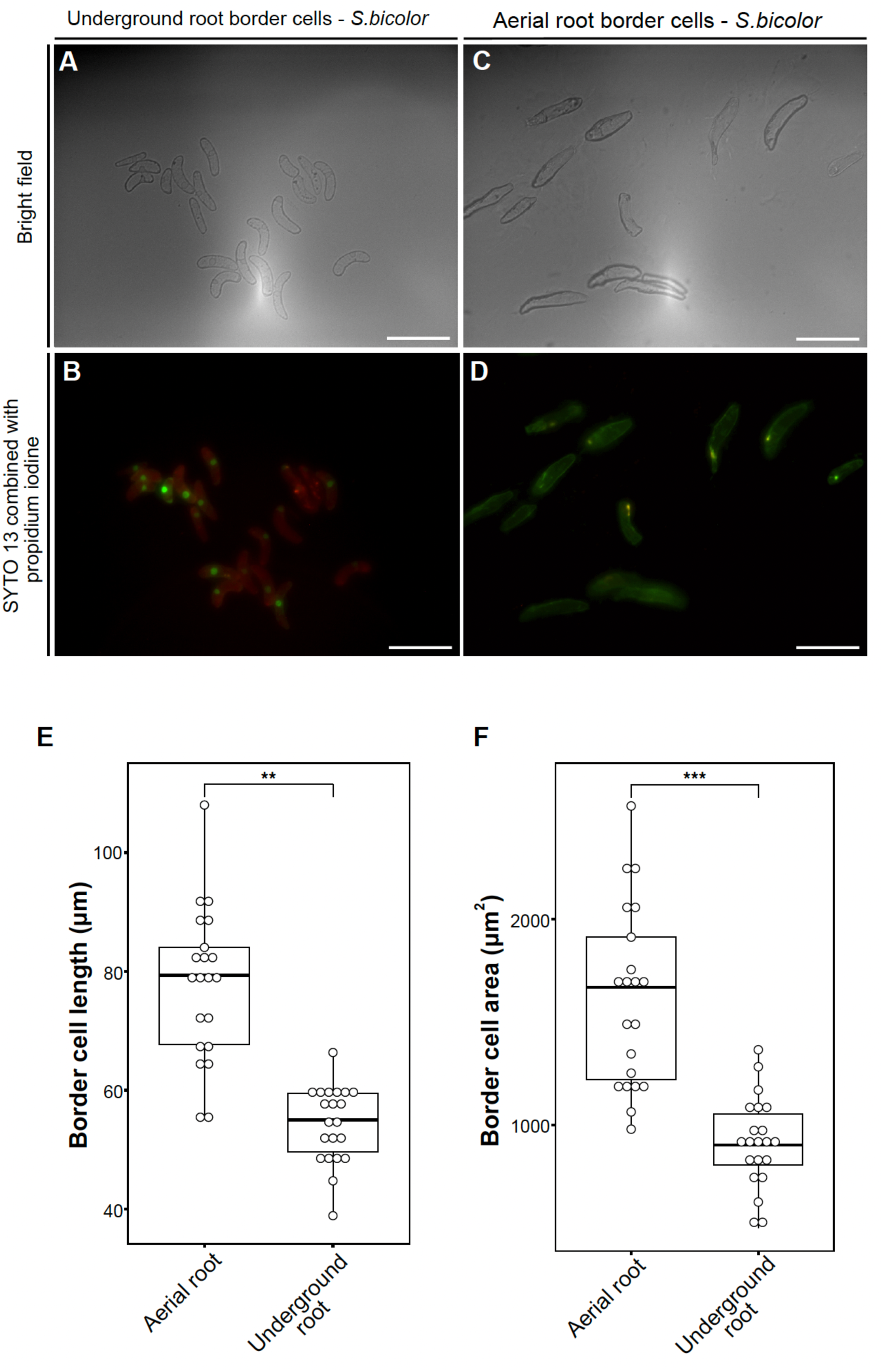
Border cells in sorghum mucilage. A) Border cells from sorghum underground roots (S=scale bar 100 μm). Border cells have a bean shape. B) Border cells from sorghum aerial roots (scale bar 100 μm). Irregular cells are observed in a more elongated form. C) Live-dead viability staining in border cells from sorghum underground roots (scale bar 100 μm). The presence of live cells was recorded using the green fluorescence of their nuclei. D) Live-dead viability staining in border cells from sorghum aerial roots (scale bar 100 μ m). Live cells were identified using the green fluorescence of their nuclei. E) Length comparison between border cells in μm. Border cells from mucilage were more elongated than border cells from underground roots. F) Area comparison between border cells in μm^2^. ** and *** denote p< 0.01 and < 0.001, respectively, based on a Wilcoxon test (n = 20).

**Suppl. Fig 4:**
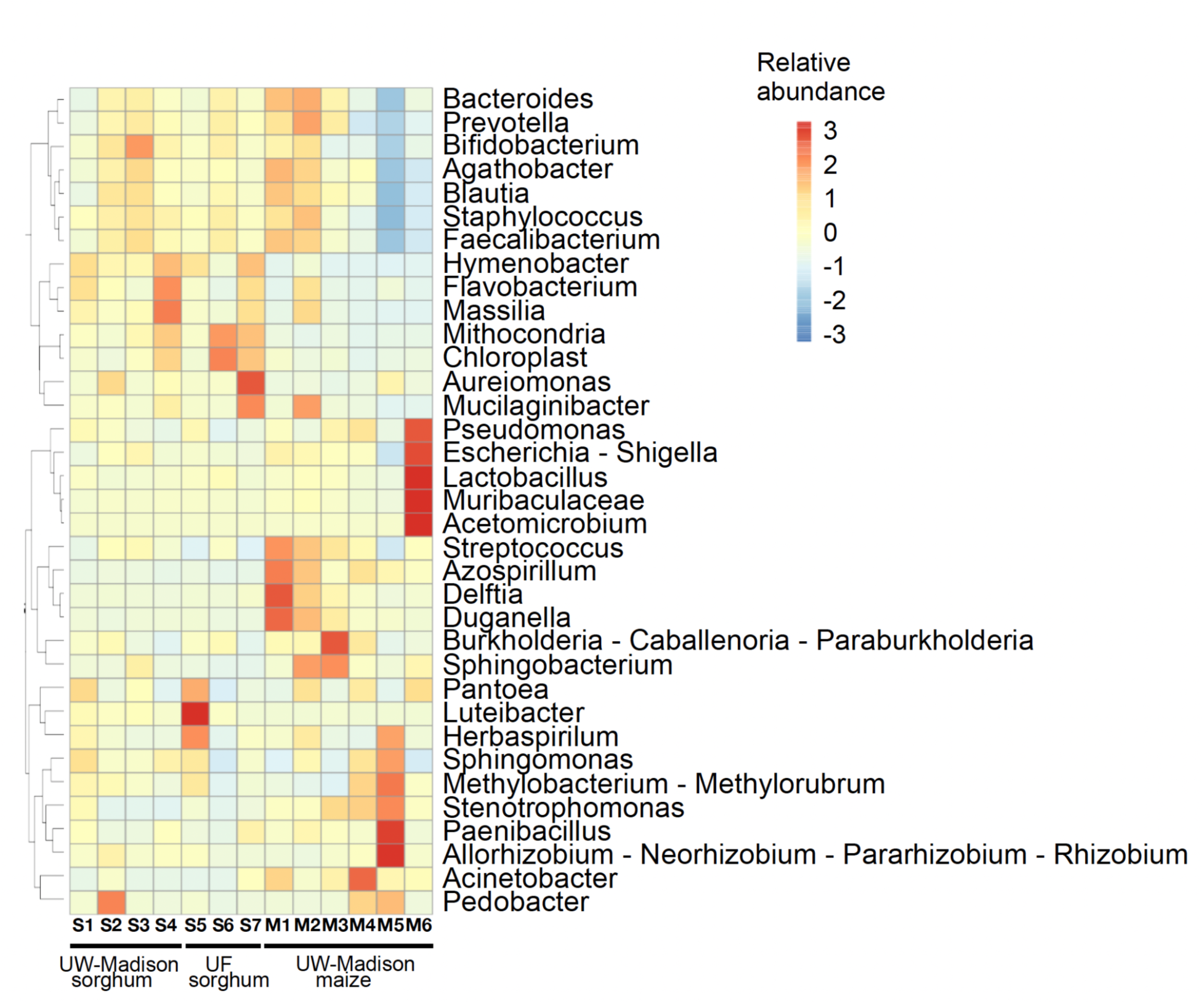
Bacterial genera from different sorghum and maize mucilage. The hierarchical clustering heatmap represents the microbiome community composition at the genus level for 13 samples obtained from the mucilage of sorghum (S) and maize (M). The samples have been grouped through clustering based on their community composition. The microbiome of each sample was obtained from different genotypes and locations, thus showing unique profiles.

**Suppl. Fig 5.**
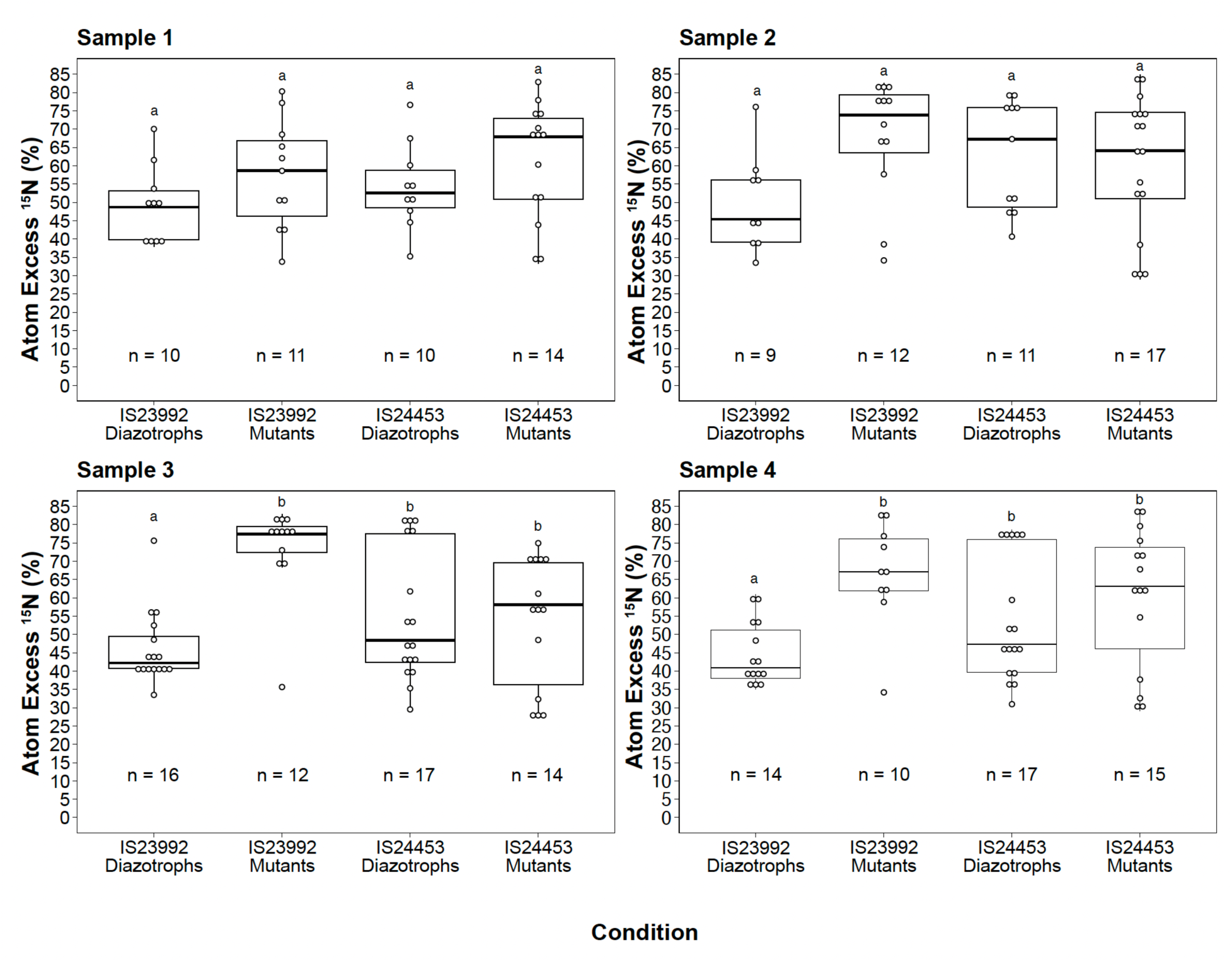
^15^N isotope dilution experiment on aerial roots of sorghum accessions from leaf samples. Accessions IS23992 and IS24453 were inoculated with the nine strains from the ARA experiment (diazotrophs), and three mutant strains *Azospirillum brasilense* FP10, *Azotobacter vinelandii* DJ100 Δ*nifH*, *Azorhizobium caulinodans* ORS571 Δ*nifA*. The third topmost leave was analyzed. The samples were collected before inoculation (sample one - stage three) and every two weeks (sample 3 to 4 - stage 4 to 6). An ANOVA was conducted using the package multcompViewto determine the factors contributing to differences in means.

**Suppl. Fig 6.**
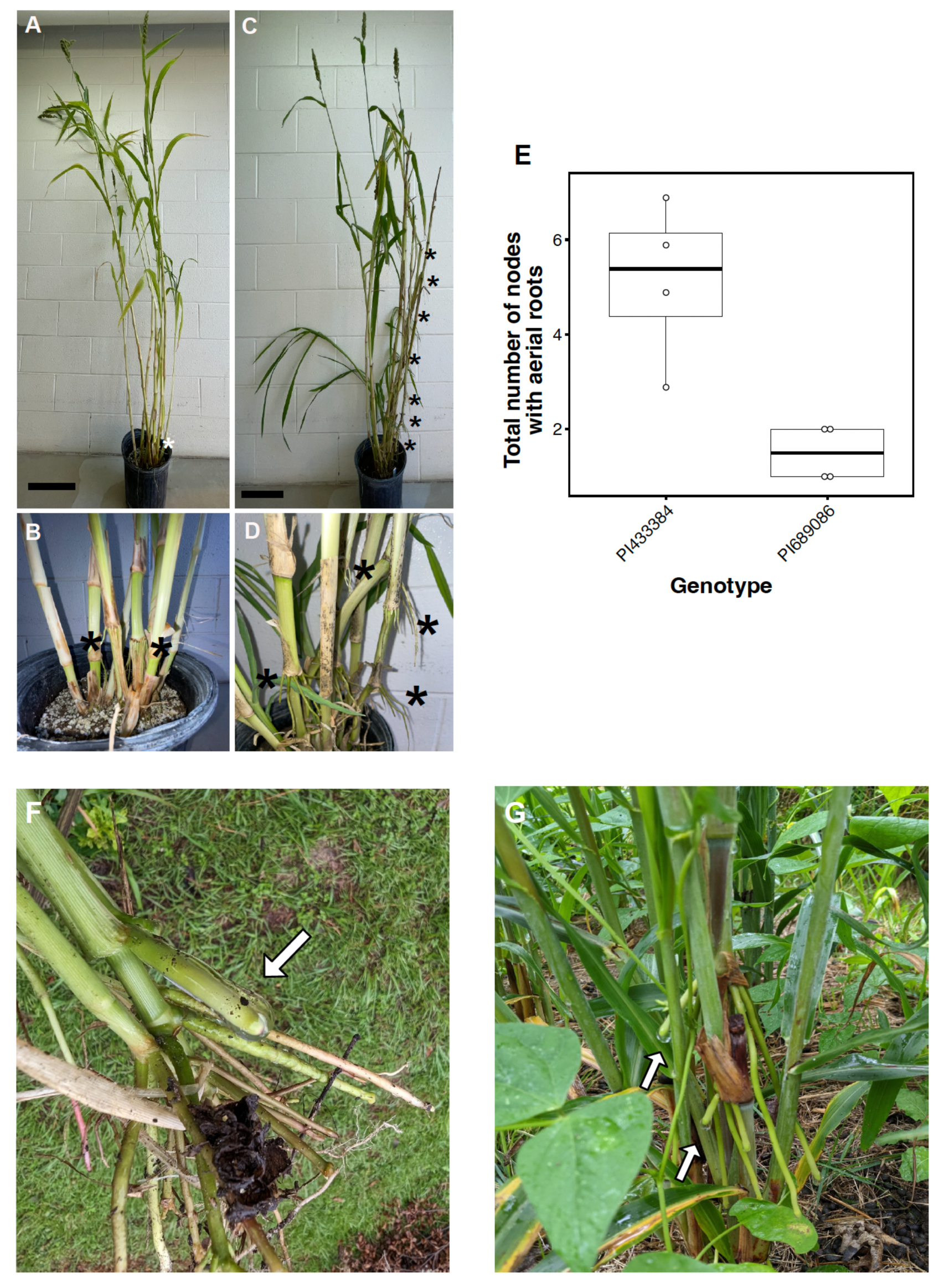
Aerial roots in other grass species. A preliminary screening in *Setaria* (*Setaria italica* and *Setaria magna*) and other species of the Sorghum clade revealed the presence of aerial roots. A) *Setaria magna* accession PI689086 displayed few aerial roots. Scale bar: 20 cm. B) Close-up of an aerial root at the base of *Setaria italica* accession PI689086. C) The *Setaria magna* accession PI433384 exhibited more aerial roots. Scale bar: 20 cm. D) Close-up of an aerial root at the base of *Setaria magna* accession PI433384. E) Quantification of aerial roots in Setaria accessions PI433384 and PI689086. F) Johnsongrass (*Sorghum halapense*) grown in Tallahassee, Florida, displayed aerial roots with mucilage. G) Sudangrass (*Sorghum bicolor* × *Sorghum arundinaceum*) grown in western North Carolina displayed aerial roots with mucilage. Asterisks mark the nodes with aerial roots; white arrows indicate aerial roots with mucilage.

**Supplementary Table 1.** Sorghum accessions to evaluate aerial root formation in greenhouse and field experiments.

| **Accession** | **Collection** | **Origin** |
| --- | --- | --- |
| IS10757 | Mini core | Chad |
| IS11026 | Mini core | Ethiopia |
| IS15170 | Mini core | Cameron |
| IS15466 | Mini core | Cameron |
| IS15478 | Mini core | Cameron |
| IS17348 | ICRISAT | Ethiopia |
| IS23992 | Mini core | Yemen |
| IS24453 | Mini core | South Africa |
| IS25089 | Mini core | Ghana |
| IS28023 | ICRISAT | Yemen |
| IS28025 | ICRISAT | Yemen |
| IS28033 | ICRISAT | Yemen |
| IS28262 | ICRISAT | Yemen |
| IS29078 | ICRISAT | Yemen |

**Supplementary Table 2.** Mucilage sugar composition of accession IS11026 in two independent experiments. Monosaccharide composition was determined by GC-MS. * data obtained from 1.64 mg sample; ** data obtained from a 2.16 mg sample; nd = not detected.

|  | **Replicate 1\*** |  | **Replicate 2\*\*** |  |
| --- | --- | --- | --- | --- |
| **Monosaccharide** | **Amount (µg)** | **% by mole** | **Amount (µg)** | **% by mole** |
| Arabinose | 163.9 | 19.77 | 254.4 | 20.66 |
| Rhamnose | nd | - | nd | - |
| Ribose | nd | - | nd | - |
| Fucose | 124.9 | 13.77 | 224.6 | 16.68 |
| Xylose | 21.1 | 2.54 | 34.1 | 2.77 |
| Glucuronic Acid | 75.9 | 7.08 | 121.8 | 7.65 |
| Galacturonic acid | nd | - | nd | - |
| Mannose | 67.8 | 6.81 | 87.1 | 5.89 |
| Galactose | 484.9 | 48.73 | 674.0 | 45.59 |
| Glucose | 12.9 | 1.29 | 11.4 | 0.77 |
| N-Acetyl Mannosamine | nd | - | nd | - |
| N-Acetyl Galactosamine | nd | - | nd | - |
| N-Acetyl Glucosamine | nd | - | nd | - |
| **Total** | **951.4** | **100** | **1,407.4** | **100** |

**Supplementary Table 3.** Mucilage samples from maize and sorghum for primary metabolism analysis.

| **Genotype** | **Plant** | **Volume (mg)** |
| --- | --- | --- |
| IS 29091 biological replicate 1 | Sorghum | 1500 |
| IS 29091 biological replicate 2 | Sorghum | 200 |
| IS 29091 biological replicate 3 | Sorghum | 1000 |
| IS 2245 biological replicate 1 | Sorghum | 150 |
| IS 2245 biological replicate 2 | Sorghum | 75 |
| IS 2245 biological replicate 3 | Sorghum | 250 |
| CIMMYT-BANK-017456 biological replicate 1 | Maize | 750 |
| CIMMYT-BANK-017456 biological replicate 2 | Maize | 750 |
| CIMMYT BANK 017456 biological replicate 3 | Maize | 500 |
| CIMMYT BANK 017456 biological replicate 4 | Maize | 500 |
| GRIN AMES 19897 biological replicate 1 | Maize | 400 |
| GRIN AMES 19897 biological replicate 2 | Maize | 500 |
| GRIN AMES 19897 biological replicate 3 | Maize | 250 |
| GRIN AMES 19897 biological replicate 4 | Maize | 100 |

**Supplementary Table 4.**
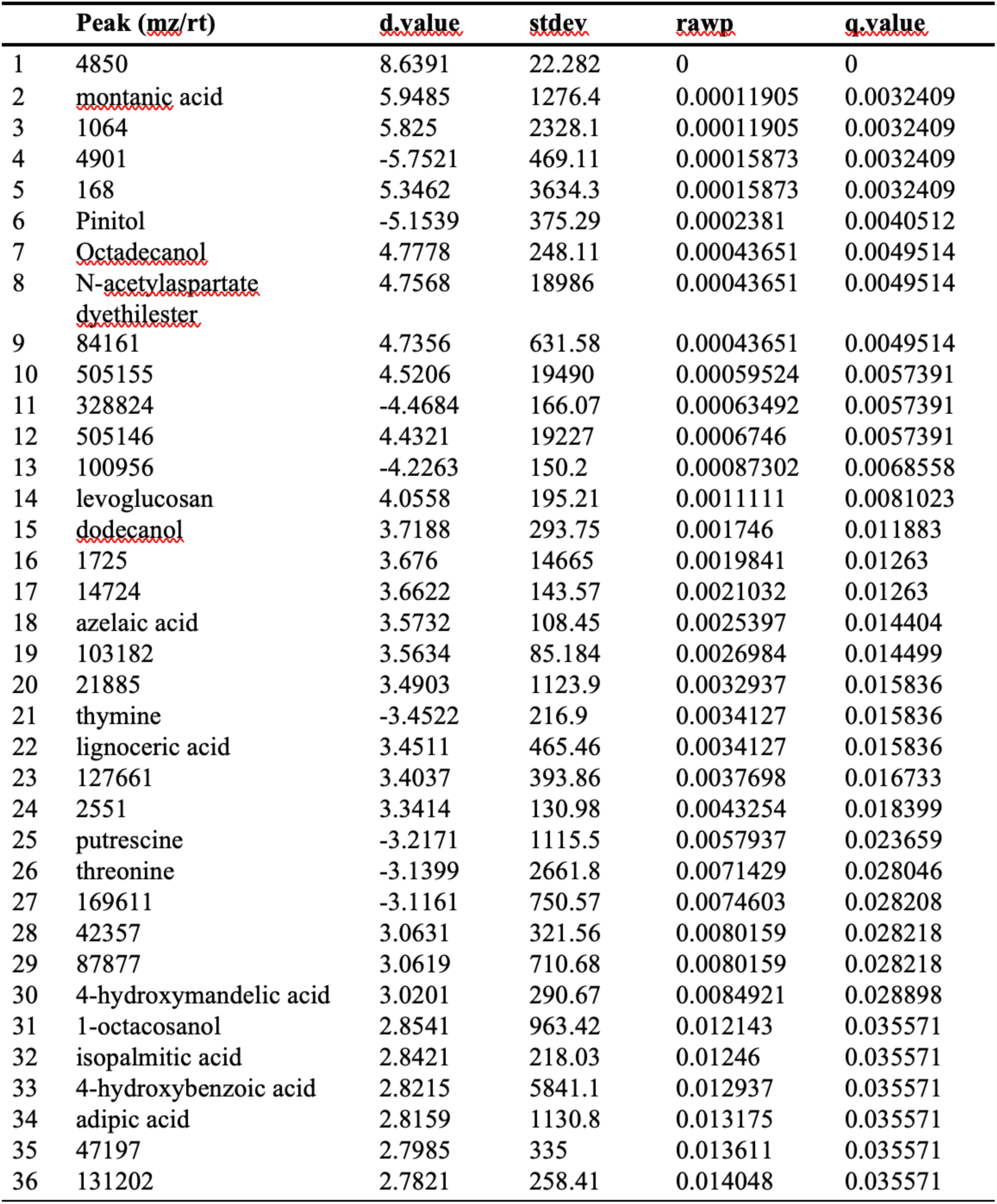
Top metabolites in aerial root mucilage. Peak (mz/rt) is the mass by charge (mz) divided by the retention time (rt). d.value denotes the comparison between sorghum and maize, where a positive value signifies that the metabolite is more abundant in sorghum than maize, and negative values indicate higher abundance in maize. Other acronyms include standard deviation (stdev), raw p-value (rawp), and q-value (q.value) for statistical analysis.

**Supplementary Table 5.** Maize and sorghum accession for microbiome study. Number of replicates is indicated in the sample ID.

| Sample ID | Genotype | Plant | Location |
| --- | --- | --- | --- |
| S1 (4 samples) | IS 2245 | sorghum | West Madison |
| S2 (3 samples) | IS 2245* | sorghum | West Madison |
| S3 (4 samples) | IS 23992 | sorghum | West Madison |
| S4 (4 samples) | IS 29092 | sorghum | West Madison |
| S5 (4 samples) | IS 2902 | sorghum | UF |
| S6 (4 samples) | IS 2902* | sorghum | UF |
| S7 (4 samples) | IS 2245 | sorghum | UF |
| M1 (4 samples) | CIMMYT-BANK-017456 | maize | West Madison |
| M2 (4 samples) | CIMMYT-BANK-017456 | maize | West Madison |
| M3 (4 samples) | CIMMYTA-BANK-014019 | maize | West Madison |
| M4 (4 samples) | GRIN AMES 19897 | maize | West Madison |
| M5 (3 samples) | GRIN AMES 19897 | maize | West Madison |
| M6 (4 samples) | GRIN AMES 19897 | maize | West Madison |

\*Samples were obtained from a low nitrogen pilot study

**Supplementary Table 6.** AMOVA for each sample comparison. S = sorghum sample, M = maize sample, and df = degrees of freedom. Statistically significant differences are indicated with asterisks: * p ≤ 0.05; ** p ≤ 0.01, and *** p ≤ 0.001

| Group | Sum of Square | df | Mean Square | Fixation Index | p-value |
| --- | --- | --- | --- | --- | --- |
| S1-M3 | 0.144504 | 1 | 0.144504 | 5.1731 | 0.005** |
| M4-M5 | 0.0855831 | 1 | 0.0855831 | 2.22751 | 0.095 |
| S2-M4 | 0.0864243 | 1 | 0.0864243 | 3.0695 | 0.038* |
| M2-M5 | 0.115087 | 1 | 0.115087 | 3.71158 | 0.029* |
| S6-S7 | 0.0614475 | 1 | 0.0614475 | 0.486006 | 0.61 |
| S3-M5 | 0.180132 | 1 | 0.180132 | 5.09055 | 0.026* |
| S1-S6 | 0.184116 | 1 | 0.184116 | 1.95325 | 0.263 |
| S5-M6 | 0.450519 | 1 | 0.450519 | 5.24901 | 0.01** |
| S3-M1 | 0.054311 | 1 | 0.054311 | 1.92444 | 0.114 |
| S2-M3 | 0.186426 | 1 | 0.186426 | 7.09958 | 0.03* |
| S2-S7 | 0.266473 | 1 | 0.266473 | 4.39073 | 0.068 |
| S4-S6 | 0.22921 | 1 | 0.22921 | 2.53538 | 0.198 |
| M4-M6 | 0.138883 | 1 | 0.138883 | 3.15016 | 0.05* |
| S2-S6 | 0.137483 | 1 | 0.137483 | 1.29883 | 0.413 |
| S5-M2 | 0.223533 | 1 | 0.223533 | 3.187 | 0.107 |
| M2-M4 | 0.0199211 | 1 | 0.0199211 | 0.70151 | 0.589 |
| M3-M5 | 0.0460803 | 1 | 0.0460803 | 1.26165 | 0.341 |
| M1-M5 | 0.1314 | 1 | 0.1314 | 3.8839 | 0.013* |
| S7-M6 | 0.740479 | 1 | 0.740479 | 10.4001 | 0.02* |
| S6-M2 | 0.225729 | 1 | 0.225729 | 2.42344 | 0.184 |
| S4-M6 | 0.183315 | 1 | 0.183315 | 5.21219 | 0.011* |
| S4-M1 | 0.0626517 | 1 | 0.0626517 | 2.86947 | 0.023* |
| S3-S4 | 0.0656918 | 1 | 0.0656918 | 2.8403 | 0.072 |
| S2-S4 | 0.0574829 | 1 | 0.0574829 | 3.29319 | 0.066 |
| S4-S7 | 0.436925 | 1 | 0.436925 | 8.27984 | 0.029 |
| S5-M3 | 0.407831 | 1 | 0.407831 | 5.45696 | 0.017 |
| S5-M1 | 0.270468 | 1 | 0.270468 | 3.73097 | 0.03* |
| M1-M2 | 0.0309953 | 1 | 0.0309953 | 1.26133 | 0.189 |
| S4-S5 | 0.231663 | 1 | 0.231663 | 3.43716 | 0.019** |
| S3-M3 | 0.16078 | 1 | 0.16078 | 5.27755 | 0.005** |
| S5-S7 | 0.0721351 | 1 | 0.0721351 | 0.697439 | 0.438 |
| S4-M2 | 0.0571673 | 1 | 0.0571673 | 2.93463 | 0.028* |
| M1-M3 | 0.115701 | 1 | 0.115701 | 3.96639 | 0.003** |
| S1-S5 | 0.163211 | 1 | 0.163211 | 2.29049 | 0.184 |
| S4-M5 | 0.0948554 | 1 | 0.0948554 | 3.42189 | 0.034* |
| S7-M5 | 0.549234 | 1 | 0.549234 | 7.74059 | 0.04* |
| S2-S5 | 0.111242 | 1 | 0.111242 | 1.4217 | 0.269 |
| S7-M4 | 0.436884 | 1 | 0.436884 | 7.08226 | 0.001*** |
| S1-M4 | 0.0736667 | 1 | 0.0736667 | 2.49594 | 0.089 |
| S7-M3 | 0.655788 | 1 | 0.655788 | 10.9105 | 0.007** |
| S3-M2 | 0.0690644 | 1 | 0.0690644 | 2.66988 | 0.034* |
| M3-M6 | 0.0679816 | 1 | 0.0679816 | 1.59931 | 0.201 |
| S3-S7 | 0.418749 | 1 | 0.418749 | 7.07855 | 0.011* |
| M5-M6 | 0.10333 | 1 | 0.10333 | 2.07341 | 0.044* |
| S2-M1 | 0.0962802 | 1 | 0.0962802 | 4.08539 | 0.032* |
| M2-M6 | 0.201935 | 1 | 0.201935 | 5.3267 | 0.014* |
| S5-M4 | 0.245334 | 1 | 0.245334 | 3.21468 | 0.023* |
| S5-M5 | 0.348851 | 1 | 0.348851 | 3.94134 | 0.027* |
| S4-M4 | 0.0506137 | 1 | 0.0506137 | 1.97263 | 0.121 |
| S6-M6 | 0.437969 | 1 | 0.437969 | 4.02419 | <0.001* |
| S4-M3 | 0.108391 | 1 | 0.108391 | 4.50182 | 0.012* |
| S2-M2 | 0.0652358 | 1 | 0.0652358 | 3.14503 | 0.025* |
| S6-M1 | 0.272552 | 1 | 0.272552 | 2.85402 | 0.048* |
| S1-S3 | 0.0566845 | 1 | 0.0566845 | 2.10059 | 0.052* |
| M1-M6 | 0.221378 | 1 | 0.221378 | 5.49822 | 0.015* |
| S1-S4 | 0.0218245 | 1 | 0.0218245 | 1.05958 | 0.349 |
| M2-M3 | 0.0991976 | 1 | 0.0991976 | 3.69911 | 0.033* |
| S2-M5 | 0.16442 | 1 | 0.16442 | 5.24333 | 0.105 |
| S1-M5 | 0.123894 | 1 | 0.123894 | 3.83004 | 0.016* |
| S6-M3 | 0.381372 | 1 | 0.381372 | 3.90187 | 0.007** |
| S1-M6 | 0.217747 | 1 | 0.217747 | 5.57941 | 0.009** |
| S2-M6 | 0.264406 | 1 | 0.264406 | 6.68187 | 0.015* |
| S3-S6 | 0.219448 | 1 | 0.219448 | 2.26721 | 0.309 |
| S1-M2 | 0.0740045 | 1 | 0.0740045 | 3.17115 | 0.014* |
| S6-M4 | 0.23653 | 1 | 0.23653 | 2.38145 | 0.211 |
| S7-M1 | 0.485294 | 1 | 0.485294 | 8.38696 | 0.007** |
| S1-S2 | 0.0290423 | 1 | 0.0290423 | 1.31515 | 0.245 |
| M1-M4 | 0.0441517 | 1 | 0.0441517 | 1.43577 | 0.138 |
| M3-M4 | 0.0719655 | 1 | 0.0719655 | 2.18114 | 0.125 |
| S1-S7 | 0.390742 | 1 | 0.390742 | 6.90035 | 0.006** |
| S5-S6 | 0.0159133 | 1 | 0.0159133 | 0.11281 | 0.878 |
| S6-M5 | 0.325414 | 1 | 0.325414 | 2.80247 | 0.157 |
| S3-S5 | 0.194684 | 1 | 0.194684 | 2.63844 | 0.07 |
| S2-S3 | 0.0518463 | 1 | 0.0518463 | 2.06391 | 0.13 |
| S3-M6 | 0.241142 | 1 | 0.241142 | 5.80253 | 0.023* |
| S1-M1 | 0.0850429 | 1 | 0.0850429 | 3.31029 | 0.016* |
| S7-M2 | 0.406276 | 1 | 0.406276 | 7.31907 | 0.032* |
| S3-M4 | 0.0837043 | 1 | 0.0837043 | 2.61203 | 0.001*** |
| All samples | 2.31435 | 12 | 0.192863 | 3.6472 | <0.001*** |

**Supplementary Table 7.**
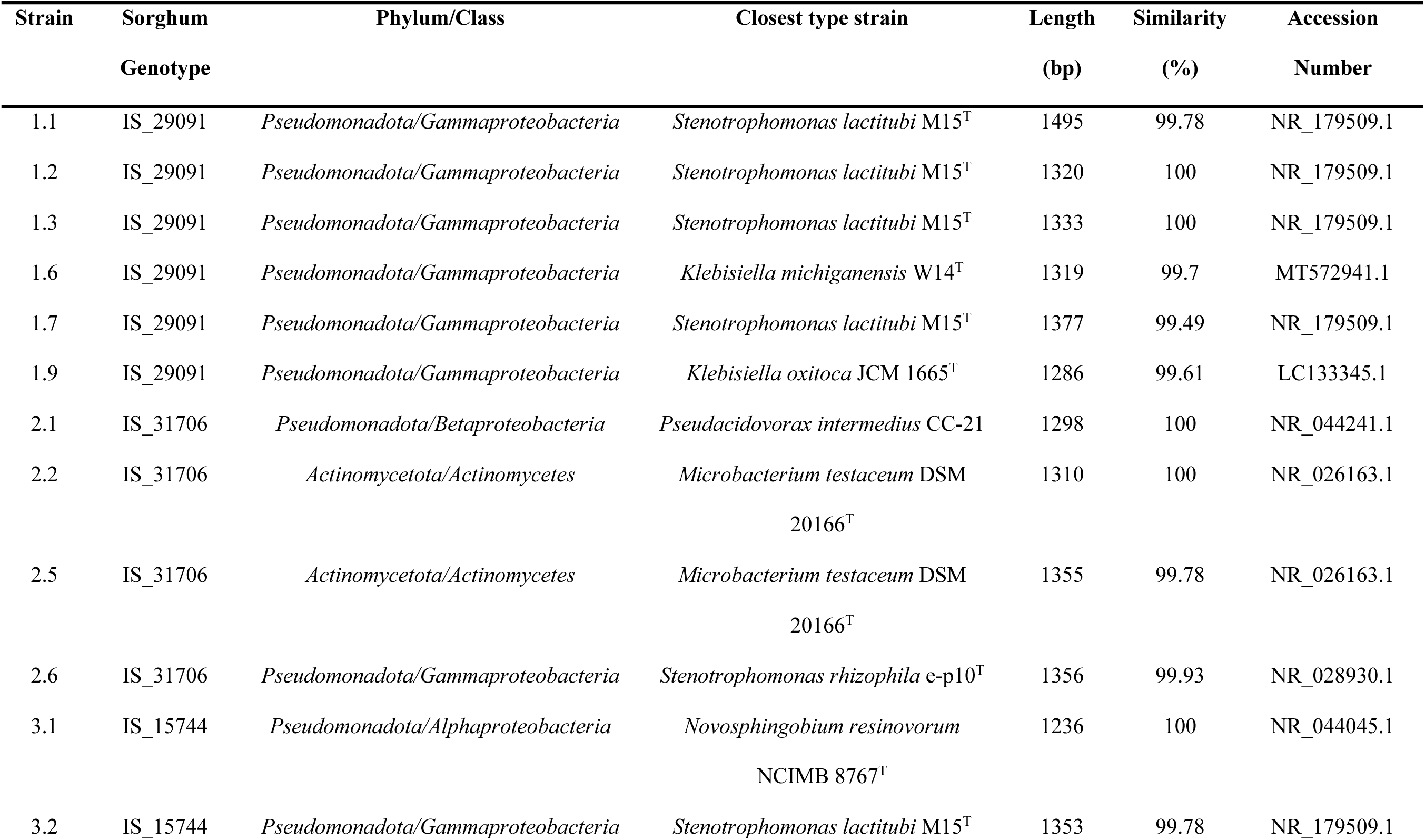
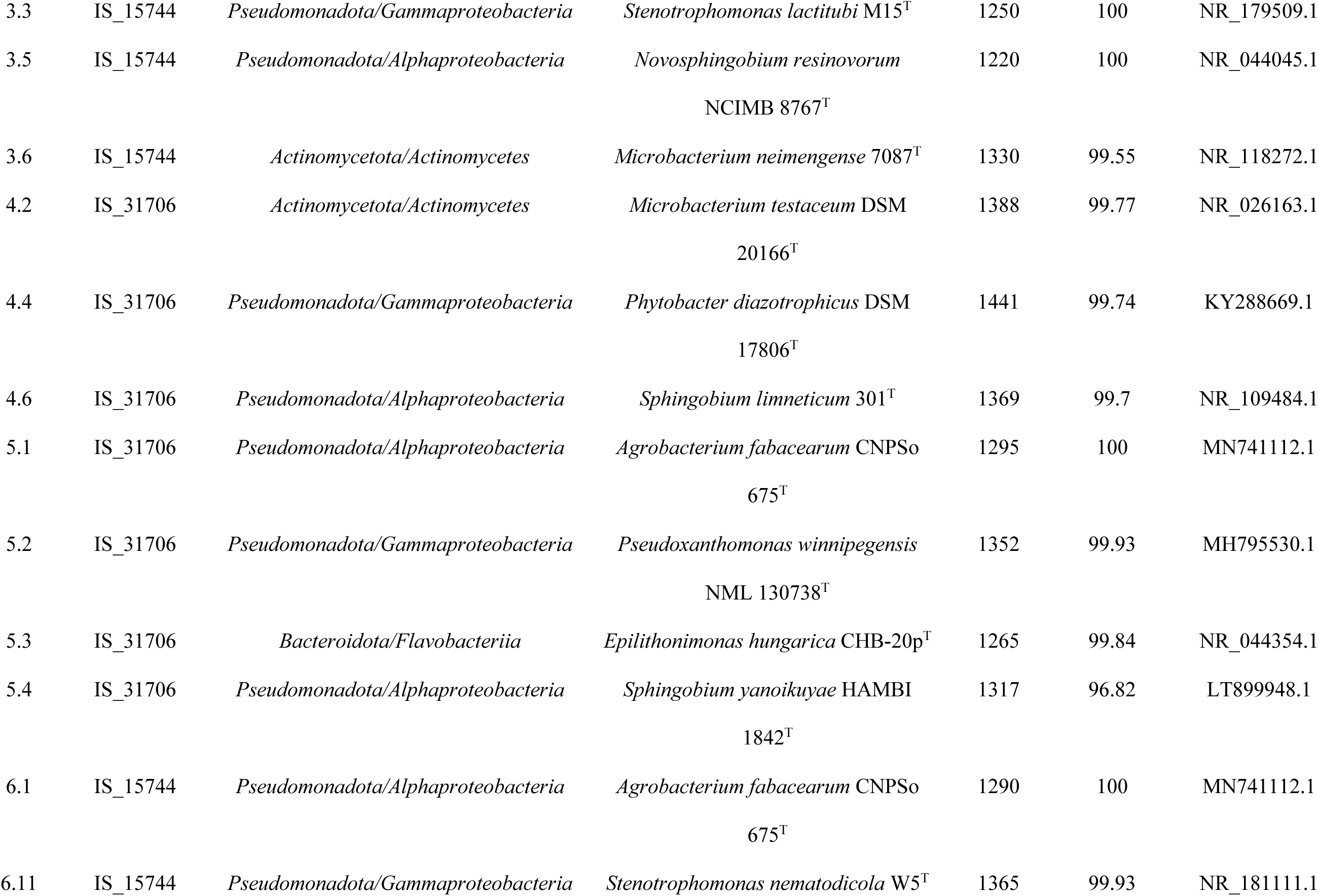
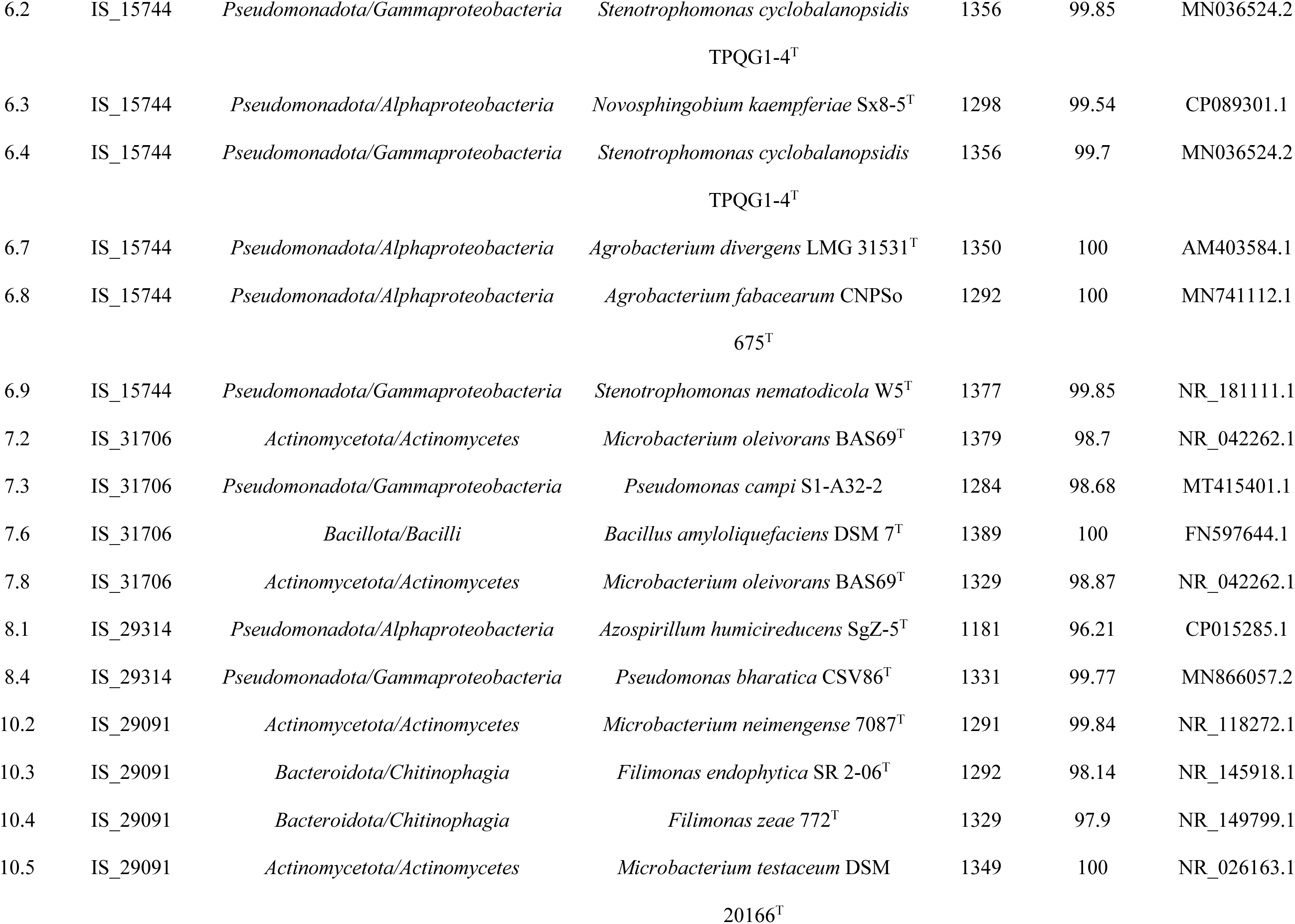
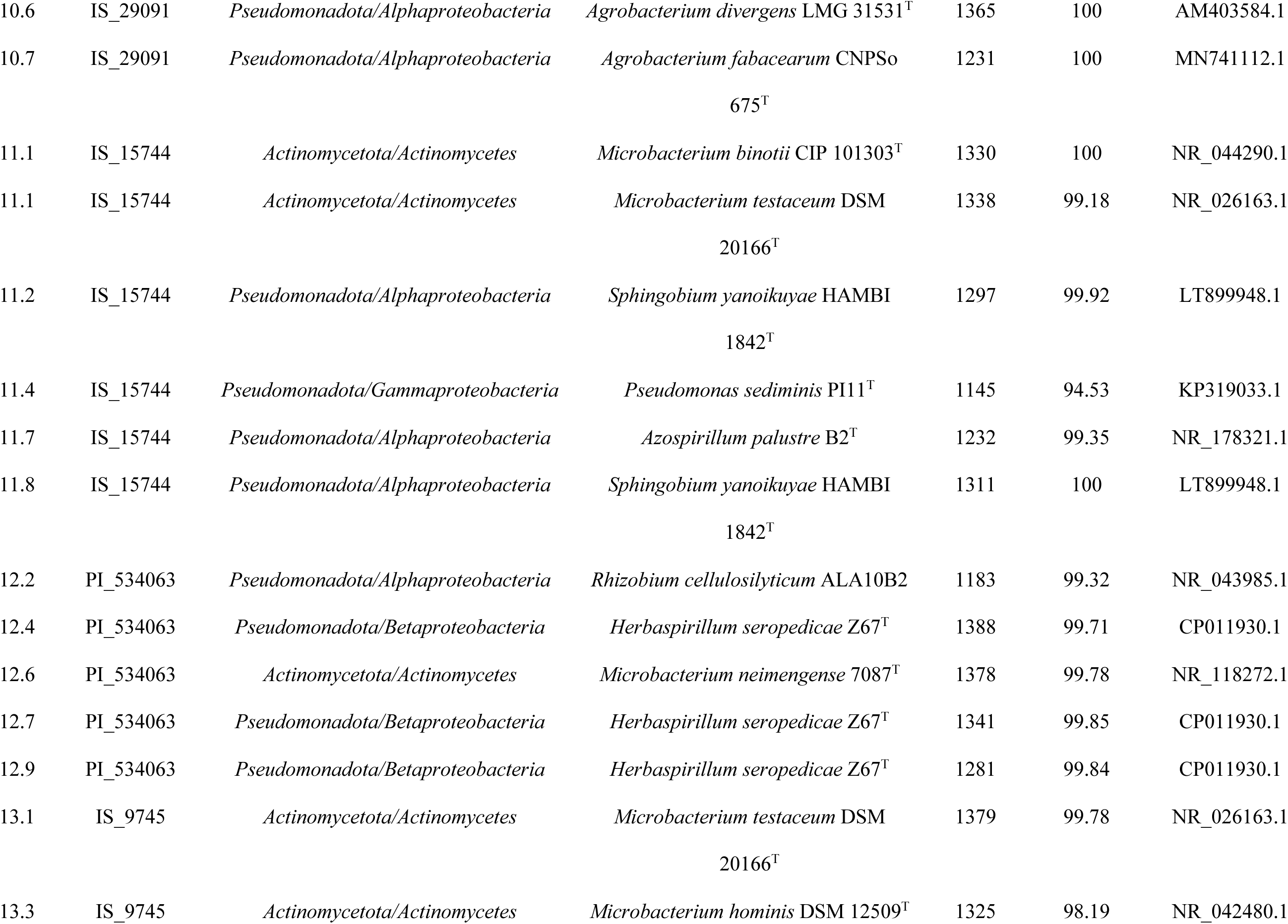
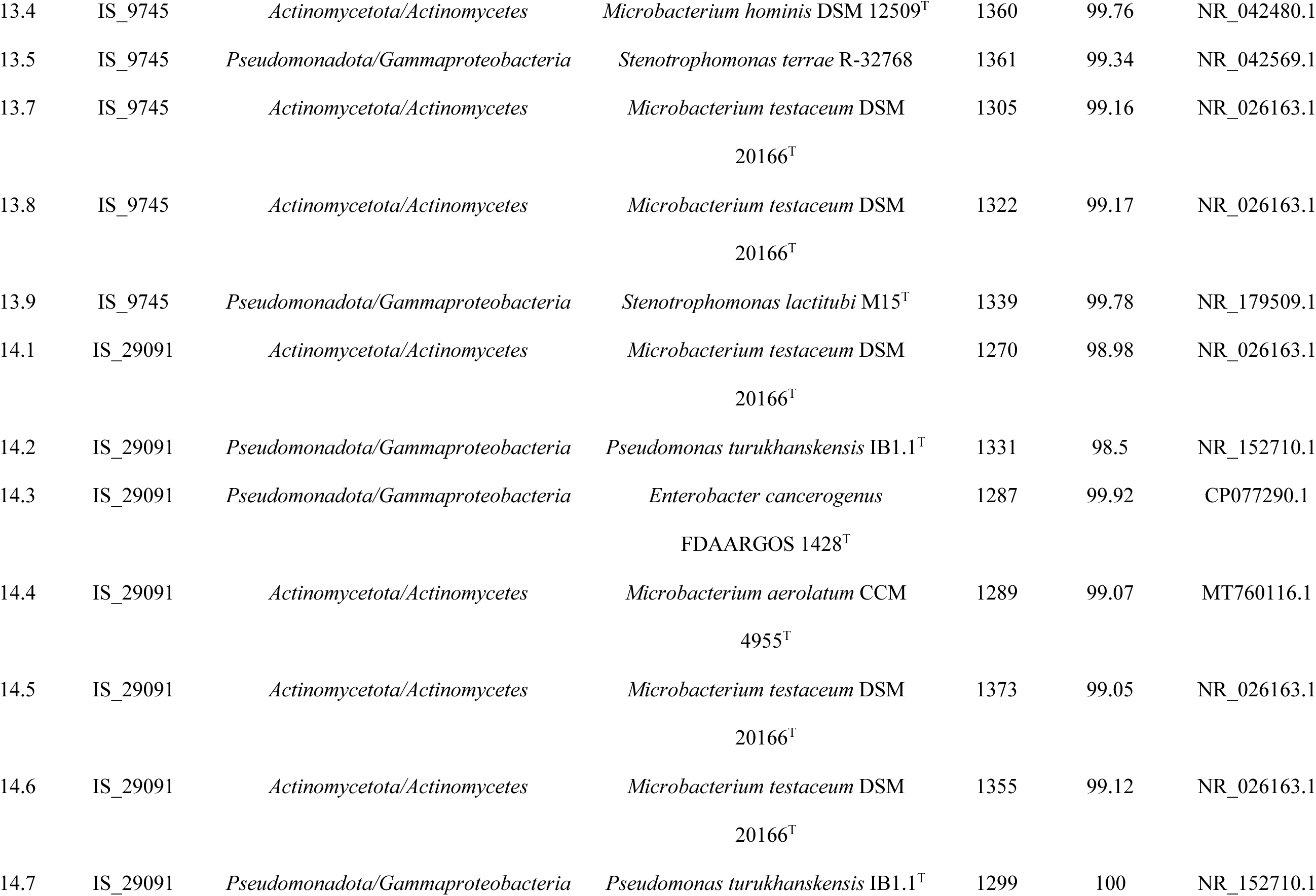
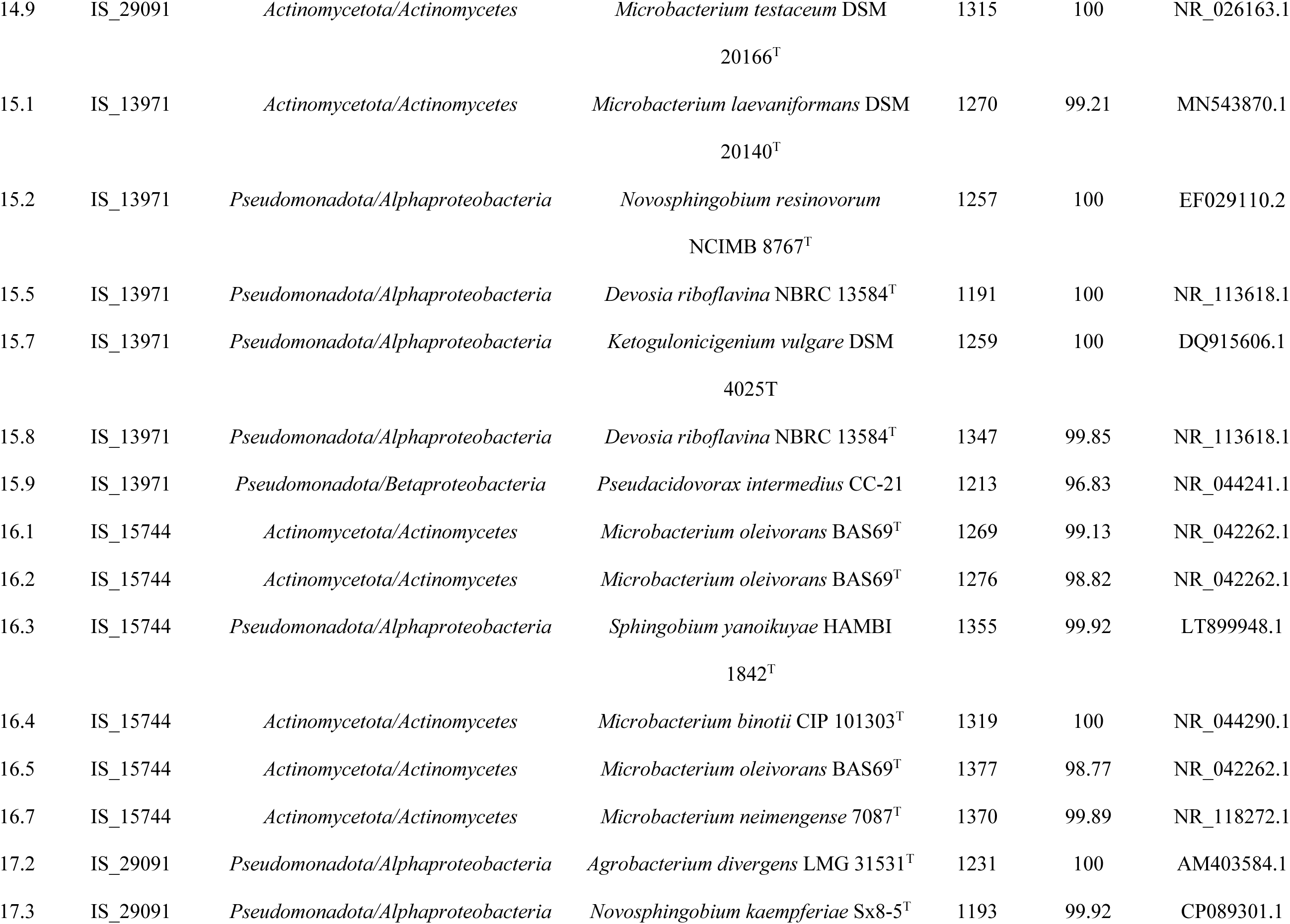
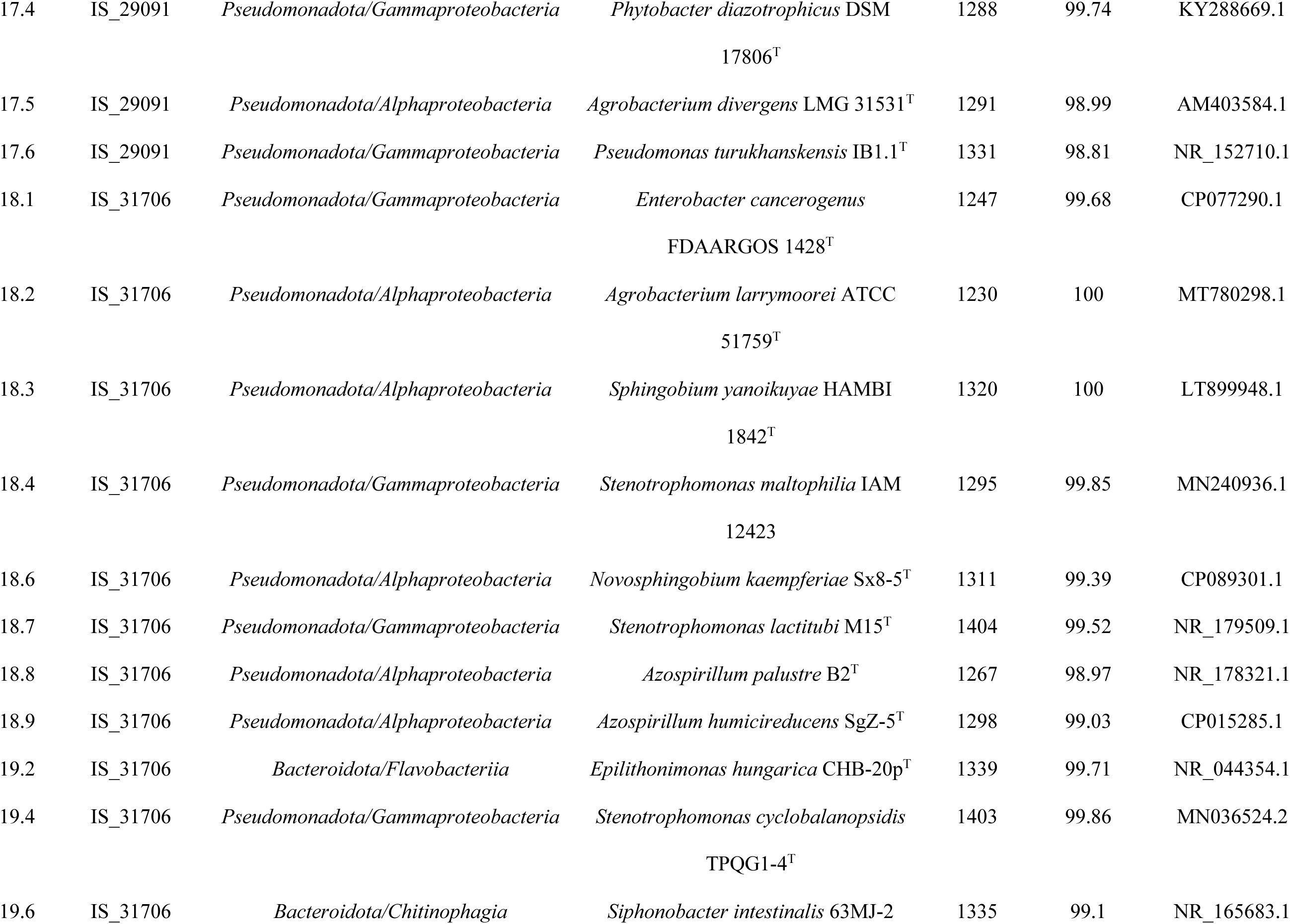
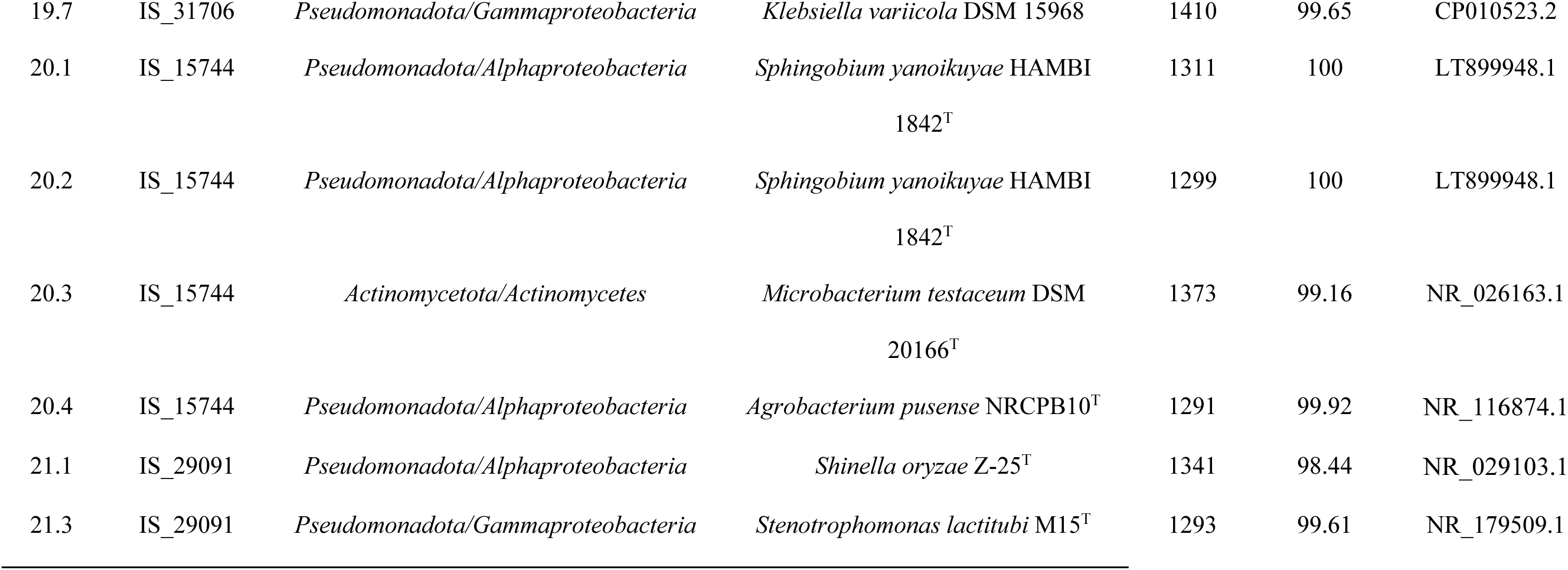
Diazotrophs isolated from the sorghum mucilage.

**Supplementary Table 8.**
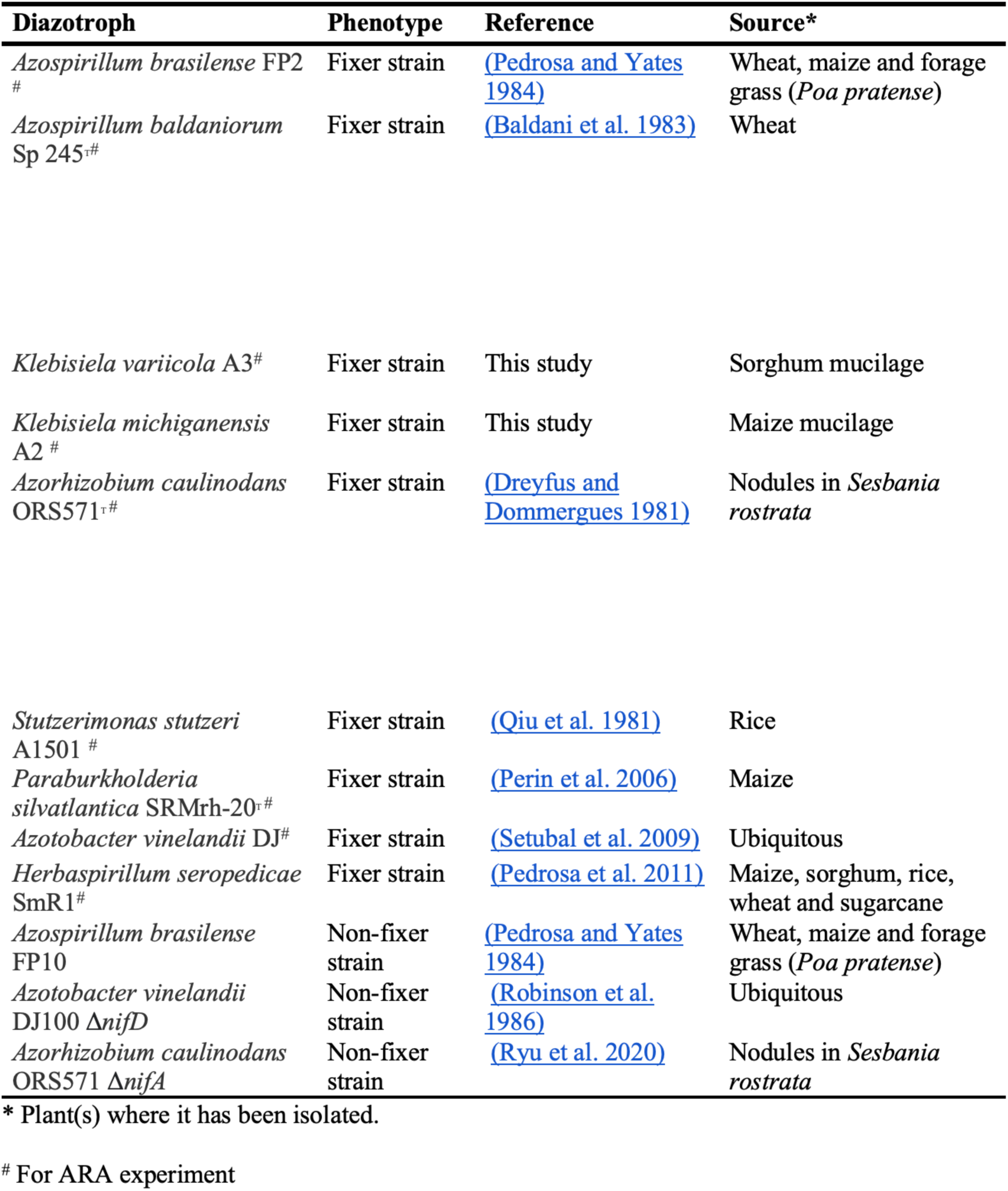
Diazotrophs that have been used for biological nitrogen fixation experiments. References to the origin of these strains and the source of isolation are provided.

